# Identification of novel buffering mechanisms in aortic arch artery development and congenital heart disease

**DOI:** 10.1101/2023.03.02.530833

**Authors:** AnnJosette Ramirez, Christina A. Vyzas, Huaning Zhao, Kevin Eng, Karl Degenhardt, Sophie Astrof

**Affiliations:** Department of Cell Biology and Molecular Medicine, New Jersey Medical School, Rutgers Biomedical and Health Sciences, Newark, NJ, 07103; Multidisciplinary Ph.D. Program in Biomedical Sciences: Cell Biology, Neuroscience and Physiology Track, New Jersey Medical School, Rutgers Biomedical and Health Sciences, Newark, NJ, 07103; Department of Statistics, Rutgers University, School of Arts and Sciences, Piscataway, NJ 08854; Children’s Hospital of Pennsylvania, University of Pennsylvania, Philadelphia, PA 19107

**Keywords:** second heart field, endothelial progenitors, pharyngeal arch arteries, aortic arch arteries, ductus arteriosus, veins, 22q11 deletion syndrome

## Abstract

**Rationale:** The resiliency of embryonic development to genetic and environmental perturbations has been long appreciated; however, little is known about the mechanisms underlying the robustness of developmental processes. Aberrations resulting in neonatal lethality are exemplified by congenital heart disease (CHD) arising from defective morphogenesis of pharyngeal arch arteries (PAA) and their derivatives.

**Objective:** To uncover mechanisms underlying the robustness of PAA morphogenesis.

**Methods and Results:** The second heart field (SHF) gives rise to the PAA endothelium. Here, we show that the number of SHF-derived ECs is regulated by *VEGFR2* and *Tbx1*. Remarkably, when SHF-derived EC number is decreased, PAA development can be rescued by the compensatory endothelium. Blocking such compensatory response leads to embryonic demise. To determine the source of compensating ECs and mechanisms regulating their recruitment, we investigated three-dimensional EC connectivity, EC fate, and gene expression. Our studies demonstrate that the expression of VEGFR2 by the SHF is required for the differentiation of SHF-derived cells into PAA ECs. The deletion of one VEGFR2 allele (VEGFR2^SHF-HET^) reduces SHF contribution to the PAA endothelium, while the deletion of both alleles (VEGFR2^SHF-KO^) abolishes it. The decrease in SHF-derived ECs in VEGFR2^SHF-HET^ and VEGFR2^SHF-KO^ embryos is complemented by the recruitment of ECs from the nearby veins. Compensatory ECs contribute to PAA derivatives, giving rise to the endothelium of the aortic arch and the ductus in VEGFR2^SHF-KO^ mutants. Blocking the compensatory response in VEGFR2^SHF-KO^ mutants results in embryonic lethality shortly after mid-gestation. The compensatory ECs are absent in *Tbx1^+/-^* embryos, a model for 22q11 deletion syndrome, leading to unpredictable arch artery morphogenesis and CHD. *Tbx1* regulates the recruitment of the compensatory endothelium in an SHF-non-cell-autonomous manner.

**Conclusions:** Our studies uncover a novel buffering mechanism underlying the resiliency of PAA development and remodeling.

**Nonstandard Abbreviations and Acronyms in Alphabetical Order:** CHD – congenital heart disease; ECs – endothelial cells; IAA-B – interrupted aortic arch type B; PAA – pharyngeal arch arteries; RERSA – retro-esophageal right subclavian artery; SHF – second heart field; VEGFR2 – Vascular endothelial growth factor receptor 2.

## Introduction

How complex developmental processes almost invariably lead to predictable phenotypes has remained a conundrum for many years. In 1942, C.H. Waddington proposed the concept of canalization of development, which refers to the tendency of developmental processes to follow a specific trajectory despite genetic and environmental perturbations ^1^. Noting that genetically “wild type” organisms have much less developmental variability than organisms with genetic mutations, Waddington argued for a buffering mechanism(s) that allow constancy in developmental outcomes despite minor genetic or environmental perturbations. One example of canalized development is the organogenesis of the cardiovascular system. The aortic arch is a major vessel that routes oxygenated blood into the systemic circulation. In the vast majority of mice and humans, this vessel forms a leftward arch to connect the heart and the left descending aorta. However, mutations affecting pharyngeal mesoderm or cardiac neural crest development can result in unpredictable defects in arch artery development and remodeling; for example, a rightward instead of a leftward arch can form, two arches can form: one rightward and one leftward, giving rise to a ring, or the aortic arch can be absent completely ^2,3^. Although rare, congenital defects offer an opportunity to understand buffering mechanisms underlying the constancy of development and how alterations in these mechanisms result in birth defects.

Congenital heart disease (CHD) is the most prevalent congenital anomaly, affecting approximately 8 individuals per 1000 live births in the United States ^4,5^. Defects in the development of pharyngeal arch artery (PAA) derivatives, the aortic arch and the ductus arteriosus give rise to lethal forms of CHD. During development, each PAA forms within one pharyngeal arch, a complex tissue in the thoracic region of the embryo containing cells derived from all three germ layers ^6^. PAAs form by vasculogenesis, where mesoderm-derived vascular progenitors populate the pharyngeal arch mesenchyme and differentiate into endothelial cells (ECs). These ECs form a primary network of small vessels which then remodel into one large artery per arch ^7–11^. We and others have shown that PAA endothelium, particularly the 4^th^ and 6^th^ PAAs, is mainly derived from a subpopulation of splanchnic mesoderm known as the Second Heart Field (SHF) ^7,8,10^.

Following their formation, PAAs undergo asymmetric remodeling to give rise to the aortic arch artery, its branches, and the ductus arteriosus ^2,3,12^. Defective formation and remodeling of the left 4^th^ PAA leads to the lethal CHD phenotype interrupted aortic arch type B (IAA-B), resulting in the loss of the lumenal continuity between the ascending and descending aorta and interrupting the circulation of oxygenated blood throughout the body ^13^. Defective formation and remodeling of the right 4^th^ PAA results in a retro-esophageal right subclavian artery (RERSA), where the right subclavian artery branches from the descending aorta. Although non-lethal, RERSA can lead to *dysphagia lusoria*, a difficulty in swallowing ^14^. The left 6^th^ PAA gives rise to the ductus arteriosus, and aberrant formation or precocious regression of this vessel leads to pulmonary atresia, which is lethal ^15,16^.

Chromosome 22q11 deletion syndrome (22q11 DS) is one of the most prevalent chromosomal microdeletion syndromes, with an incidence of 1 in 3,000-6,000 live births ^17,18^. Characterized by multiple anomalies, 80% of 22q11DS cases present with CHD ^18,19^. Approximately 50-60% of patients with interrupted aortic arch have 22q11DS, with IAA-B being the most prevalent subtype ^20–23^. *Tbx1* is among the genes deleted in 22q11DS, and knockout studies in mice demonstrated that heterozygosity of *Tbx1* results in aberrant 4^th^ PAA development, phenocopying aortic arch defects in 22q11DS patients ^24–26^.

When investigating the impact of global *Tbx1* heterozygosity on the development of the fourth PAAs, we observed that aberrant 4^th^ PAA development in *Tbx1^+/-^* embryos was accompanied by a decrease in the number of SHF-derived ECs in the 4^th^ pharyngeal arches. A similar reduction in SHF-derived EC numbers was noted when a single copy of the VEGFR2 gene was deleted in the SHF. However, unlike *Tbx1^+/-^* embryos, embryos with a heterozygous deletion of VEGFR2 in the SHF contained compensatory endothelium that mitigated the decreased numbers of SHF-derived ECs and rescued arch artery development. In this manuscript, we describe the discovery of this compensatory endothelium, identify its source, its role in arch artery remodeling, and the importance of VEGFR2 and Tbx1 in its regulation. Collectively, this work provides valuable insights into the resilience of developmental mechanisms, shedding light on the robustness and plasticity involved in the development of PAA-derived vasculature.

## Results

### The expression of VEGFR2 in the SHF is required for the specification of pharyngeal arch artery endothelium

To determine whether the SHF is a principal source of the PAA endothelium, we ablated VEGFR2 from the SHF using the *Isl1^Cre^* knock-in strain by crossing *VEGFR2^+/-^;Isl1^Cre/+^* males with *VEGFR2^f/f^; Rosa^mTmG/mTmG^*females, generating *VEGFR2^flox/-^; Rosa^mTmG/+^; Isl1^Cre/+^* (*VEGFR2^Isl^*^1^*^-KO^*), *VEGFR2^flox/+^; Rosa^mTmG/+^; Isl1^Cre/+^* (*VEGFR2^Isl^*^1^*^-Het^*), and *VEGFR2^flox/+^; Rosa^mTmG/+^; Isl1^+/+^* (*VEGFR2^Isl^*^1^*^-WT^*) embryos. Embryos were examined at embryonic day (E) 10.5 when PAA formation was nearly complete (34 – 36s) ^7,8,27^. Vasculature was visualized using whole-mount immunofluorescent staining (WM-IF) and antibodies against VEGFR2, followed by confocal imaging of the entire pharyngeal region and three-dimensional (3D) reconstruction of optical sections. We hypothesized that deleting VEGFR2 from the SHF would abolish the PAA endothelium. Our analyses showed that PAA formation in *VEGFR2^Isl^*^1^*^-Het^* and *VEGFR2^Isl^*^1^*^-KO^* embryos was defective, compared with *VEGFR2^Isl^*^1^*^-WT^*(**Fig. S1A-F)**; the 6^th^ PAA of *VEGFR2^Isl^*^1^*^-Het^* embryos was hypoplastic (compare **Fig. S1A-B** with **Fig. S1C-D**), whereas the 4^th^ and 6^th^ PAAs of *VEGFR2^Isl^*^1^*^-KO^*embryos were either hypoplastic or aplastic (**Fig. S1E-F, Table S1**). Surprisingly, despite the deletion of VEGFR2 in the SHF (**Fig. S1G-I, outlined**), VEGFR2+ endothelium was present in all three pharyngeal arches of *VEGFR2^Isl^*^1^*^-KO^* embryos (**Fig. S1E-F**). These results suggested that either VEGFR2 expression was not essential for the differentiation of SHF-derived vascular progenitors into PAA ECs or that mutant embryos contained ECs from a source other than the SHF.

To test these hypotheses and determine the fate of SHF-derived mesoderm in mutant embryos, we used the *Mef2C-AHF-Cre* transgenic strain and *Rosa^tdTom/tdTom^* reporter. For these experiments, *VEGFR2^+/-^; Mef2C-AHF-Cre* males were crossed with *VEGFR2^flox/flox^*; *Rosa^tdTom/tdTom^* females to generate *VEGFR2^flox/-^; Mef2C-AHF-Cre+; Rosa^tdTom/+^* (*VEGFR2^SHF-KO^*) embryos and *VEGFR2^flox/+^; Mef2C-AHF-Cre+; Rosa^tdTom/+^* (*VEGFR2^SHF-Het^*) embryos. *VEGFR2^+/-^; Mef2C-AHF-Cre* males were also crossed with *Rosa^tdTom/tdTom^* females to generate *VEGFR2^+/+^; Mef2C-AHF-Cre+; Rosa^tdTom/+^* (*VEGFR2^SHF-WT^*) embryos. To verify the ablation of VEGFR2 in the *Mef2C-AHF*-lineage, we performed WM-IF and used antibodies against VEGFR2 and tdTomato, encoded by the Cre-Reporter transgene. In *VEGFR2^SHF-WT^* embryos, Cre-reporter+ cells in the anterior and posterior poles of the dorsal pericardial wall and pharyngeal arches expressed VEGFR2 (**Fig. 1A-I**). In *VEGFR2^SHF-KO^* embryos, VEGFR2 expression was absent in the dorsal pericardial wall (**Fig. 1J-N, outlined**), and 84-88% of Cre-reporter+ cells in the pharyngeal arches did not express VEGFR2 (**Figs. 1O-R arrows, Table S2**). These experiments demonstrated the successful ablation of VEGFR2 protein from the *Mef2c-AHF*-lineage in *VEGFR2^SHF-KO^* embryos.

**Figure 1.**
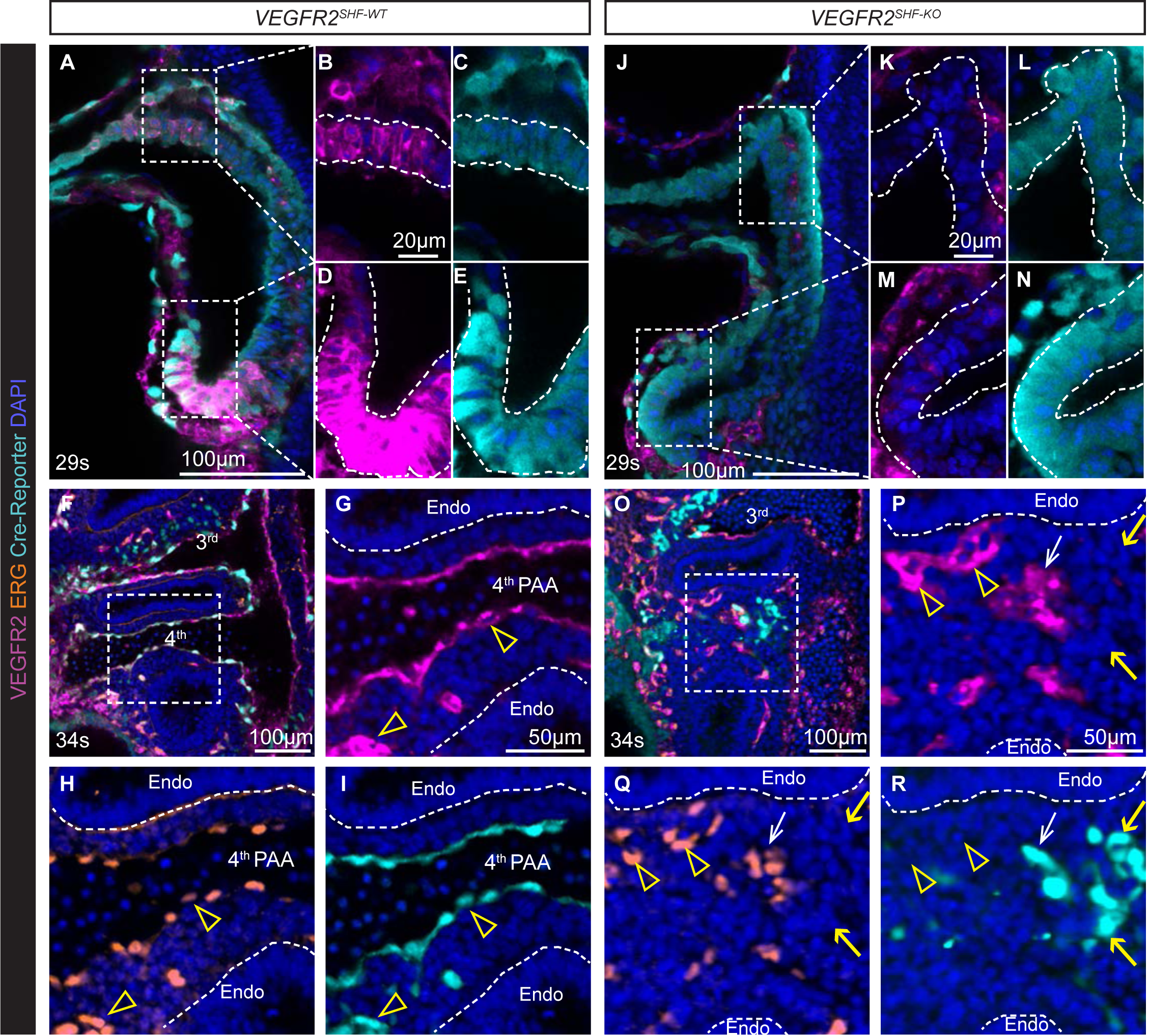
Ablation of VEGFR2 in the second heart field (SHF)-derived cells using *Mef2c-AHF-Cre* transgenic strain. Embryos were stained using whole mount immunofluorescence to detect *Mef2c-AHF-Cre* lineage cells (cyan), VEGFR2 (magenta), and ERG (orange). Optical 0.45 μm-thick confocal sections are shown. **A-E,** E10.0 *VEGFR2^SHF-WT^* embryos express VEGFR2 in the anterior, **B-C** and posterior dorsal pericardial wall **D-E**, outlined. Magnifications in **B-E** are the same. **F-I,** 4th pharyngeal arch ECs in E10.5 *VEGFR2^SHF-WT^* embryos are derived from the SHF. The box in **F** is placed in the 4th arch and is expanded in **(G-I)**. Arrowheads point to VEGFR2+ (magenta), ERG+ (orange), and a Cre-reporter+ (cyan) ECs in the 4th pharyngeal arch. **J-N,** VEGFR2 is downregulated in E10 *VEGFR2^SHF-KO^*embryos in the anterior, **K-L,** and posterior dorsal pericardial wall, **M-N**, outlined. Magnifications in **K-N** are the same. **O-R,** Most *VEGFR2^SHF-KO^* embryos lack a central PAA in the 4th arch and 4th pharyngeal arch ECs in E10.5 *VEGFR2^SHF-KO^* embryos are derived from a non-SHF source. The box in **O** is placed in the 4th pharyngeal arch and is expanded in **P-R**. Arrowheads point to Cre-reporter negative, ERG+ ECs. Yellow arrows point to Cre-reporter+ VEGFR2-negative SHF-derived cells; White arrow points to an escapee Cre-reporter+VEGFR2+ EC. Endoderm (endo) is outlined by dashed lines in **G-I** and **P-R** to demonstrate the equivalent anatomical positions of the 4th arch of *VEGFR2^SHF-WT^*and *VEGFR2^SHF-KO^* embryos.

To determine whether the expression of VEGFR2 in the *Mef2c-AHF*-lineage was required for PAA development, we examined PAA formation in *VEGFR2^SHF-WT^*, *VEGFR2^SHF-Het^*, and *VEGFR2^SHF-KO^* embryos at E10.5. Embryos were stained by WM-IF using antibodies against VEGFR2, the ETS transcription factor ERG, and the Cre-reporter tdTomato. As with the *Isl1^Cre/+^*strain, we observed defective PAA formation in *VEGFR2^SHF-Het^* and *VEGFR2^SHF-KO^* embryos (**Fig. 2A-F, Table S1**). *VEGFR2^SHF-Het^*embryos had hypoplastic 6^th^ PAAs (**Fig. 2C-D**). PAA defects in *VEGFR2^SHF-KO^*mutants were more severe than in the *VEGFR2^SHF-Het^*: most 4^th^ and 6^th^ PAAs were aplastic, that is, the pharyngeal endothelium remained in the plexus form without forming a central artery (**Fig. 2E-F**).

**Figure 2.**
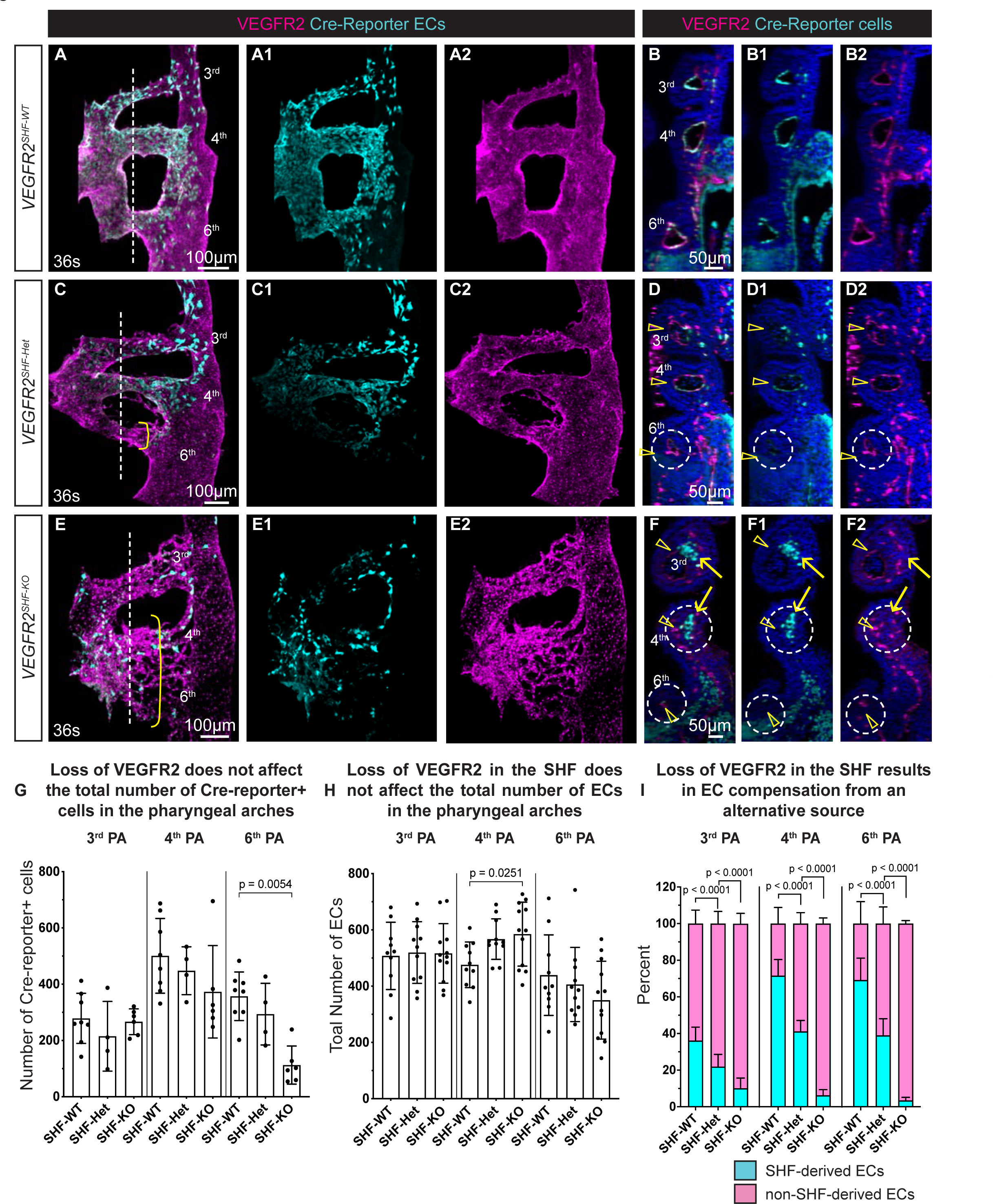
Expression of VEGFR2 is required for SHF-derived cells to give rise to pharyngeal arch artery ECs. Whole mount immunofluorescence and confocal imaging of E10.5 embryos to detect VEGFR2 (magenta), Cre-reporter (cyan) and nuclei (blue). **A-A2, C-C2, E-E2,** Imaris was used to segment vasculature and generate 3D maximum intensity projections. Pharyngeal vasculature and the dorsal aorta were segmented, and only Cre-reporter+ cells within the segmented vessels are shown. **A-A2,** *VEGFR2^SHF-WT^*, **C-C2,** *VEGFR2^SHF-Het^*, and **E-E2,** *VEGFR2^SHF-KO^*embryos. Yellow brackets highlight aberrant PAAs in **C, E**. Dashed lines mark the levels of coronal optical sections through the pharyngeal arches shown in **B, D,** and **F**. Dotted circles mark aberrant PAAs in *VEGFR2^SHF-Het^* and *VEGFR2^SHF-KO^* embryos. Arrowheads in **D-D2** and **F-F2** point to non-SHF-derived cells. Arrows in **F-F2** point to VEGFR2-negative SHF-derived cells. **G,** Quantification of SHF-derived cells in each pharyngeal arch. *VEGFR2^SHF-WT^* n = 10 arches, *VEGFR2^SHF-Het^*n = 4 arches, *VEGFR2^SHF-KO^* n = 6 arches. **H,** Quantification of total EC numbers in each pharyngeal arch. *VEGFR2^SHF-WT^*n = 10 arches, *VEGFR2^SHF-Het^* n = 12 arches, *VEGFR2^SHF-KO^*n = 12 arches. Each dot marks one pharyngeal arch. Bars show standard deviation. p values were calculated using 2-tailed, unpaired, non-parametric Kruskal-Wallis tests with Dunn’s correction for multiple testing; only p<0.05 are displayed. **I,** The proportion of Cre-reporter+ ECs and Cre-reporter-negative ECs in each pharyngeal arch. Means ± SD are shown. *VEGFR2^SHF-WT^*n = 8 arches, *VEGFR2^SHF-Het^* n = 11 arches, *VEGFR2^SHF-KO^*n = 11 arches. Statistical differences in proportions of SHF- and non- SHF-derived cells were determined, as described in Methods.

To quantify the total number of SHF-derived cells, total EC number, and the number of SHF-derived EC in the pharyngeal arches, we used the Imaris software to segment the pharyngeal arches as well as the endothelium within the arches, as described ^28^. Although the total numbers of SHF-derived cells (**Table S2**) and ECs (**Table S3**) were similar across all the genotypes (**Fig. 2G-H**), the proportion of pharyngeal arch ECs derived from the SHF was decreased in *VEGFR2^SHF-Het^*and *VEGFR2^SHF-KO^* embryos (compare **Fig. 2E** and **2C** with **2A**, wherein only Cre-reporter+ ECs are shown, quantified in **Fig. 2I**). Compared with controls, the proportion of SHF-derived ECs was reduced about 2-fold in *VEGFR2^SHF-Het^* (blue bars in **Fig. 2I**). These data indicated that the expression of one copy of VEGFR2 was not sufficient to generate wild-type levels of SHF-derived ECs.

In *VEGFR2^SHF-KO^* embryos, the proportions of SHF-derived ECs were decreased by 3-, 10-, and 20-fold compared with controls in the 3^rd^, 4^th^, and 6^th^ pharyngeal arches, respectively (cyan bars in **Fig. 2I, Table S3**). The striking paucity of SHF-derived endothelium in the arches of the mutants is seen in **Fig. 2**; Compare the abundance of cyan ECs in control (**Fig. 2A-A1)** with that in *VEGFR2^SHF-KO^* **(Fig. 2E-E1)**. Pharyngeal arch histology is shown in coronal optical sections through the arches (**Fig. 2B-B2, D-D2, F-F2**). SHF-derived cells are mainly found in the endothelium in control and *VEGFR2^SHF-Het^* embryos (**Fig. 2B-B2, D-D2**); whereas, in *VEGFR2^SHF-KO^* embryos, most SHF-derived cells lacked VEGFR2 and were not incorporated into the endothelium (**Fig. 2F-F2**, arrows).

Among pharyngeal arch ECs in *VEGFR2^SHF-KO^* embryos, a small minority (6.3%, or ∼37 cells) were SHF-derived (**Fig. 2I**, **Table S4**). However, most of these SHF-derived ECs in *VEGFR2^SHF-KO^* embryos still expressed VEGFR2 (e.g., white arrow in **Fig. 1P-R** and **Table S4**); we termed these cells escapees. In these cells, Cre-mediated recombination occurred between the loxP sites in the ROSA locus but not the VEGFR2 locus, leading to the expression of both the fluorescent reporter and VEGFR2. Alternatively, Cre-mediated recombination at the VEGFR2 locus in these cells had not yet resulted in decreased VEGFR2 protein levels by the time of their differentiation into ECs. The remaining 12% of SHF-derived ECs in the 4^th^ pharyngeal arch of *VEGFR2^SHF-KO^* embryos (∼4 cells) lacked VEGFR2 (**Table S4**). To summarize, about 4 out of a total of 585 ECs in the 4^th^ arch ECs of *VEGFR2^SHF-KO^* embryos lacked VEGR2, indicating the essential role and a stringent requirement for VEGFR2 expression in the differentiation of SHF-derived progenitors into arterial endothelium and contribution to the pharyngeal arch arteries. Finally, similar phenotypes resulted from VEGFR2 deletion using either the *Isl1^Cre^* or *Mef2C-AHF-Cre* strains (**Fig. S1** and **Tables S5-S6**), indicating that the expression of *VEGFR2* in the SHF, the common Cre expression domain between these two strains, regulates the differentiation of SHF-derived vascular progenitors into pharyngeal arch ECs.

### Compensatory endothelium contributes to PAA-derived vasculature

Defective PAA formation leads to abnormal morphogenesis of their derivatives, the ductus arteriosus and the aortic arch and its branches ^29^. Since the total number of pharyngeal arch ECs was similar among all genotypes (**Fig. 2H**), our experiments show that an alternative vascular source compensates for the reduction in SHF-derived ECs in *VEGFR2^SHF-Het^* and *VEGFR2^SHF-KO^* embryos to bring up the numbers of ECs in pharyngeal arches to control levels. In the 4^th^ pharyngeal arch, the total EC number in *VEGFR2^SHF-KO^*embryos was slightly higher than in controls (**Fig. 2H**, **Table S3**), likely due to the increased proliferation of the compensatory, non-SHF ECs (discussed below).

To determine if the altered balance in the numbers of SHF- and non-SHF-derived progenitors of the pharyngeal arch endothelium in *VEGFR2^SHF-Het^*and *VEGFR2^SHF-KO^* embryos influenced the remodeling of PAAs into their adult-like configuration, we examined embryos heterozygous or null for VEGFR2 in either the *Mef2C-AHF-Cre* or *Isl1^Cre^* lineages at or after E14.5, when PAA remodeling is complete. A Mendelian distribution of each genotype was observed between E14.5 and P0, indicating that the loss of SHF-derived vascular progenitors does not cause prenatal lethality (**Table S7**). To examine the morphogenesis of the aortic arch, branches, and ductus, embryos were analyzed at E14.5 or E18.5 by micro-computed tomography or conventional histology and H&E staining. On average ∼83% of embryos null for VEGFR2 in the SHF from either the *Isl1^Cre^*or *Mef2C-AHF-Cre* lines developed variable defects in PAA-derived vessels (**Fig. 3A-G1; Table S5**). The left 4^th^ PAA gives rise to the aortic arch artery segment between the left carotid artery and the left subclavian artery, and defective development of this artery leads to interrupted aortic arch type B (IAA-B) ^2,3^. Defects in morphogenesis of the vessels derived from the left or the right 4^th^ PAA were observed in 17 of 23 conditional VEGFR2-null embryos (**Table S5**). This indicates that compared with the vessels arising from the 3^rd^ and 6^th^ PAAs, morphogenesis of the vessels derived from the 4^th^ PAAs is more sensitive to the decreased contribution of SHF-derived ECs.

**Figure 3.**
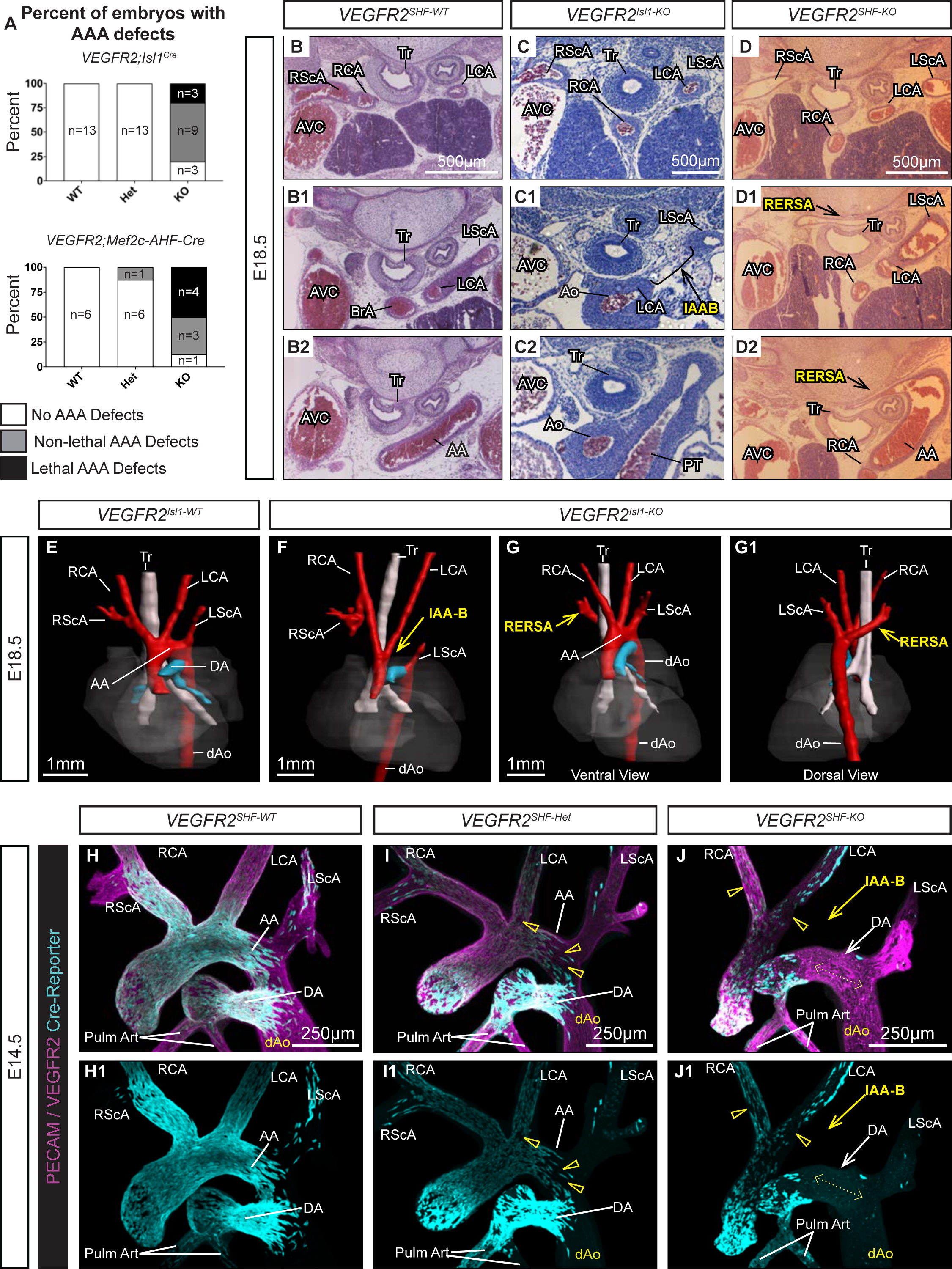
SHF-derived endothelial cells ensure consistent arch artery morphogenesis. **A,** Percent of *VEGFR2;Isl1^Cre^*(*VEGFR2^Isl^*^1^*^-WT^, VEGFR2^Isl^*^1^*^-Het^,* and *VEGFR2^Isl^*^1^*^-KO^*) and *VEGFR2;Mef2c-AHF-Cre* (*VEGFR2^SHF-WT^, VEGFR2^SHF-Het^,* and *VEGFR2^SHF-KO^*) embryos without AAA defects (white), with non-lethal AAA defects (gray), or lethal AAA defects (black). n = the number of embryos with each phenotype. **B-D2,** Transverse serial sections of E18.5 embryos stained with hematoxylin and eosin. **B-B2,** *VEGFR2^SHF-WT^* embryos. **C-C2,** IAA-B and **(D-D2)** RERSA in *VEGFR2^Isl^*^1^*^-KO^* and *VEGFR2^SHF-KO^*embryos, respectively. **E-G1,** Micro-computed tomography of E18.5 *VEGFR2^Isl^*^1^*^-WT^,* **E** and *VEGFR2^Isl^*^1^*^-KO^*, **F-G1** embryos. Arrows point to examples of IAA-B, **F** and aberrant RScA, **G-G1** in yellow. **H-J1,** 900-micron-thick vibratome sections were stained to detect ECs (purple) and SHF-derived cells. 3D reconstructions and maximum intensity projections are shown. Angled views from the dorsal sides are shown. **H-H1,** *VEGFR2^SHF-WT^* embryos. Most ECs of the carotids, the aortic arch, ductus, pulmonary trunk and ascending aorta are SHF-derived (cyan). **I-I1,** *VEGFR2^SHF-Het^* embryos. Arrowheads point to patches of non-SHF-derived ECs. Note depletion of SHF-derived cells in the aortic arch compared with *VEGFR2^SHF-WT^* embryos. **J-J1,** Large patches of non-SHF- derived cells (arrowheads) are seen in *VEGFR2^SHF-KO^* embryos. The entire DA, marked by the double-headed dashed arrow is devoid of SHF-derived cells. The intensity of cyan channel in **J1** was increased to show background fluorescence making it easier to visualize the DA. **Abbreviations:** AA - aortic arch, Ao - Ascending aorta, AVC - anterior vena cava, BrA – brachiocephalic artery; dAo - descending aorta, DA – ductus arteriosus; IAA-B – interrupted aortic arch type B; LCA and RCA – left and right carotid artery; LScA and RScA – left and right subclavian artery; PT – pulmonary trunk; Pulm Art - pulmonary arteries; RERSA – retroesophageal right subclavian artery; TR – trachea.

Despite a significantly decreased number of SHF-derived ECs in the PAAs, only 1 of 21 (4.8%) embryos heterozygous for VEGFR2 in the SHF had aberrant patterning of PAA-derived vasculature (**Table S6**). In this embryo, the right subclavian artery, which normally arises from the right 4^th^ PAA, was absent, and instead, the subclavian artery formed at the dorsal side of the descending aorta, resulting in the retroesophageal right subclavian artery, RERSA (**Fig. S2A- A2, Table S6,** embryo 21). These data suggest that the presence of a sizable proportion of SHF-derived ECs in the 4^th^ PAAs, ∼40% on average in *VEGFR2^SHF-Het^*embryos (**Fig. 2I**), ensures the constancy of PAA morphogenesis into the expected mature arrangement of the aortic arch and its branches, and the ductus arteriosus.

Intriguingly, the majority of *VEGFR2^SHF-Het^* and *VEGFR2^SHF-KO^* embryos were viable despite the decreased contribution of SHF-derived cells to the PAA endothelium of *VEGFR2^SHF-Het^* embryos and the near complete absence of SHF-derived ECs from the 4^th^ pharyngeal arches in *VEGFR2^SHF-KO^* embryos (**Table S5**). To visualize the cellular composition of PAA-derived ECs in 3D and determine whether compensatory ECs contributed to the PAA-derived vasculature, we dissected embryos at E14.5 when PAA remodeling was complete and generated 900-micron-thick vibratome sections which were then stained to detect SHF-derived cells and ECs (**Fig. 3H-J1**). In addition, we obtained thin histological sections to visualize EC composition at a higher resolution (**Fig. S2B-E3**). In controls, the majority of cells in the aortic arch, the carotids, and the ductus arteriosus were composed of SHF-derived ECs (**Fig. 3H-H1**), as we previously reported ^7^. PAA-derived vessels in *VEGFR2^SHF-Het^* embryos contained patches of non-SHF-derived endothelium, particularly noticeable in the aortic arch and the carotids (**Fig. 3I-I1**, arrowheads). Two *VEGFR2^SHF-KO^* embryos analyzed using this method had IAA-B (**Fig. 3J-J1**); However, the ductus in these mutants was entirely formed from the compensatory endothelium (**Fig.3J-J1**, dashed double-headed arrow), and the carotids contained large patches of non-SHF-derived ECs (**Fig. 3J-J1**, arrowhead).

To examine the cellular composition of PAA-derived vessels at a higher resolution, we stained tissue sections from *VEGFR2^SHF-WT^, VEGFR2^SHF-Het^,* and *VEGFR2^SHF-KO^* embryos. These experiments confirmed that PAA-derived vasculature in controls is largely composed of SHF-derived ECs (**Fig. S2B-B3**). In contrast, patches of compensatory ECs were present in PAA-derived blood vessels of *VEGFR2^SHF-Het^* embryos (**Fig. S2C-C3**), while the aortic arch and ductus of *VEGFR2^SHF-KO^* embryos were mainly derived from compensatory ECs (**Fig. S2D-D3, S2E-E3**). Together, these experiments show that compensatory ECs can give rise to large segments of PAA-derived vasculature when the SHF-derived ECs are depleted.

### Downregulation of VEGFR2 in the SHF leads to ectopic venous sprouting as well as venous gene expression and increased proliferation of compensatory ECs

To gain insights into the cellular origin of the compensatory endothelium, we dissected embryos at earlier developmental time points. Whole mount staining, confocal imaging, and 3D reconstructions of the entire pharyngeal region showed a general paucity of endothelia in the pharyngeal arches of 20-somite *VEGFR2^SHF-KO^* embryos compared with controls (**Fig. 4A-C**, blue arrows). We also noticed that the cardinal and pharyngeal veins (CV and PV) in *VEGFR2^SHF-Het^* and *VEGFR2^SHF-KO^* embryos, unlike *VEGFR2^SHF-WT^* controls, had prominent sprouts extending toward and into the pharyngeal arches (**Fig. 4B-B1, 4C-C1,** red and yellow arrows, yellow arrowheads). To better visualize vascular connections between the arterial and venous vessels, we used Imaris to segment vessels with lumenal connections to the dorsal aorta in pink and vessels with lumenal connections to CV and PV in cyan (**Fig. 4A1-C1**). These analyses were complemented by serial optical sections (**Fig. 4A2-A3, B2-B3,** and **C2-C3**). In controls, the endothelium of the pharyngeal arches had luminal continuity with the dorsal aorta (**Fig. 4A1**), and venous sprouts present at this time did not have lumenal connections with the pharyngeal vasculature (**Fig. 4A1**, and optical sections in **Fig. 4A2-A3**). In contrast, prominent venous sprouts extended from the CV and plexiform PV into the pharyngeal arches of *VEGFR2^SHF-Het^* and *VEGFR2^SHF-KO^* embryos and were lumenally contiguous with pharyngeal arch endothelium (**Fig. 4B-B3, 4C-C3**). At 27-29 somites, a large fraction of 4^th^ pharyngeal arch endothelium in *VEGFR2^SHF-KO^* embryos expressed venous markers *Apj* (**Fig. 4D-E1, 4F**) and COUPTF2 (**Fig. 4G-H1, 4I**).

**Figure 4.**
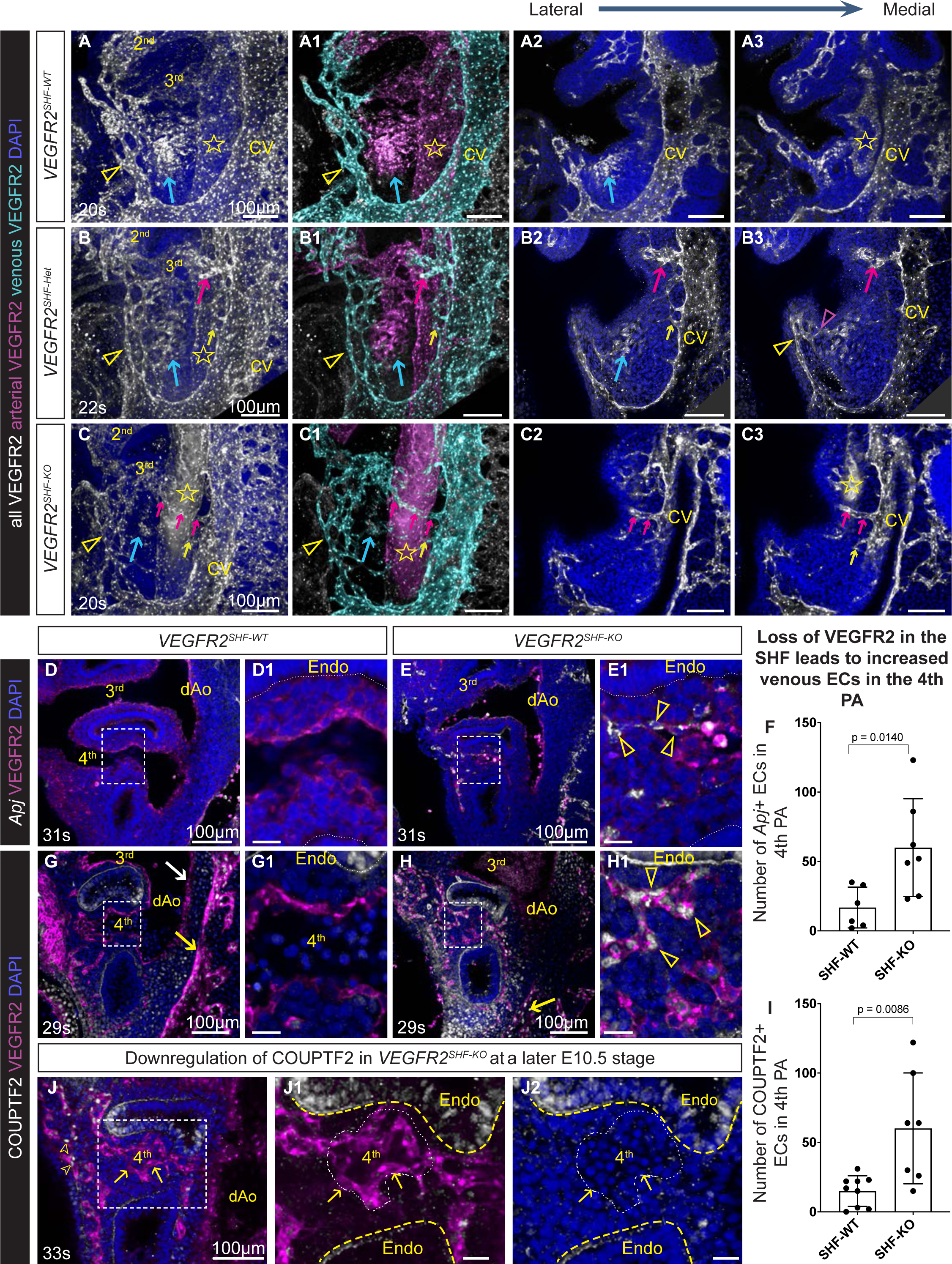
Downregulation of VEGFR2 in the SHF results in ectopic sprouts from veins and venous gene expression in the pharyngeal region. Whole mount immunofluorescence for VEGFR2 at E9.5. Stars in all panels mark the dorsal aorta, CV-cardinal vein. **A-C,** 3D reconstruction of the entire VEGFR2+ vasculature on the right side. 2^nd^ and 3^rd^ pharyngeal arches are marked. In all panels, blue arrows point to the vascular plexus in the future pharyngeal arches 4 and 6. Arrowheads point to the pharyngeal vein (PV) plexus. Images in **A-C** were modified using Imaris in **A1-C1** to mark vessels having lumenal continuity with the cardinal vein (CV) in cyan, and vessels with lumenal continuity with the dorsal aorta (marked by a star) in magenta. **A2-A3, B2-B3, C2-C3,** 10 μm serial optical sections to show venous sprouts. Arrows of the same color follow the same vessels in each row. Note sprouts from the CV and PV (arrowheads) in *VEGFR2^SHF-Het^* and *VEGFR2^SHF-KO^* embryos. Scale bars in **A-C3** are 100 μm. **D-J2**, Confocal, 0.45 μm optical slices from E10.0 embryos stained by immunofluorescence (IF) to detect VEGFR2 protein, by *in situ* hybridization to detect *Apj mRNA* in **D-E1**, or IF for VEGFR2 and COUPTF2 proteins in **G-H1, J-J2**. Boxes around the 4th arches are expanded to the right. Arrowheads point to *Apj+* 4th arch ECs in *VEGFR2^SHF-KO^* mutant in **E1** and to COUPTF2+ 4th arch ECs in **H1**. **F,** Total number of *Apj*+ ECs in the 4th arches. **I,** Total number of COUPTF2+ ECs in the 4th arches. Each dot marks one arch, p values were calculated using 2-tailed, unpaired, non-parametric Mann-Whitney tests. **J-J2**. COUPTF2 is downregulated in the 4th pharyngeal arch ECs at E10.5 in *VEGFR2^SHF-KO^* embryos. Nascent 4th PAA is outlined in **J1-J2**, arrows point to ECs. Endoderm (endo) is marked by dotted lines. Scale bars in **D1, E1, G1, H1, J1, J2** are 25 m.

Total EC numbers in pharyngeal arches of *VEGFR2^SHF-KO^*embryos become comparable with EC numbers in controls by E10.5 (**Fig. 2H**), implying robust proliferation of compensatory ECs after their arrival. To test this hypothesis, we analyzed EdU incorporation into the 4^th^ pharyngeal arch endothelium of *VEGFR2^SHF-WT^* controls and *VEGFR2^SHF-KO^* mutants. The majority (94 + 3 %) of ECs in the 4^th^ pharyngeal arches of *VEGFR2^SHF-KO^* mutants are of <u>non</u>-SHF origin (**Fig. 2I, Table S3**). EdU incorporation into the 4^th^ pharyngeal arch ECs of *VEGFR2^SHF-KO^* mutants was ∼two-fold higher than in *VEGFR2^SHF-WT^* controls, where 72 ± 9 % of 4^th^ arch ECs are SHF-derived (**Fig. S2F-H**). These data show that compensatory ECs proliferate more robustly than 4^th^ pharyngeal arch ECs in *VEGFR2^SHF-WT^* controls, most of which are SHF-derived. This finding is consistent with the observation that compensatory ECs give rise to large segments of PAA-derived vasculature at E14.5 in *VEGFR2^SHF-KO^* mutants (**Fig. 3J-J1, Fig. S2D-D3, S2E-E3**).

We found that eventually, the venous marker COUPTF2 was downregulated in 4^th^ pharyngeal arch ECs that have formed a PAA lumen in *VEGFR2^SHF-KO^* embryos (**Fig. 4J-J2**) and at E14.5 compensatory endothelium in *VEGFR2^SHF-Het^*and *VEGFR2^SHF-KO^* embryos expressed arterial marker Sox17, as in controls (**Fig. S2B3-E3**). These experiments suggested that the compensatory endothelium in the pharyngeal arches arose via angiogenesis from the nearby veins and that compensatory ECs proliferate, reorganize into a larger lumenized vessel, downregulate venous markers, and become arterialized.

### *Tbx1* regulates the abundance of SHF-derived ECs and the recruitment of the compensatory endothelium in a cell-type-specific manner

Lethal defects arising from aberrant development of the left 4^th^ PAA, such as IAA-B, are observed in about 19% of patients with 22q11DS ^19,30^. These patients have one functional copy of the *Tbx1* gene. To investigate the role of *Tbx1* in PAA development and the contribution of SHF-derived progenitors to the PAA endothelium, we crossed *Tbx1^+/-^; Mef2C-AHF-Cre* males with *Rosa^tdTom/tdTom^* females. *Tbx1^+/-^* (*Tbx1^+/-^; Mef2C-AHF-Cre)* and control (*Tbx1^+/+^; Mef2C-AHF-Cre*) embryos were collected at E10.5, and the total EC number and the number of SHF-derived ECs were quantified in the 4^th^ pharyngeal arches. Consistent with prior reports ^31^, we found that the 4^th^ PAAs in *Tbx1^+/-^* embryos were either hypoplastic or aplastic (**Fig. 5A-A5, Table S1**). Quantification of EC populations in the 4^th^ pharyngeal arches showed that unlike in *VEGFR2^SHF-Het^* embryos, the total EC number in the 4^th^ pharyngeal arches of *Tbx1^+/-^* mice was reduced by ∼33% compared with controls (**Fig. 5D**, compare the first two bars).

**Figure 5.**
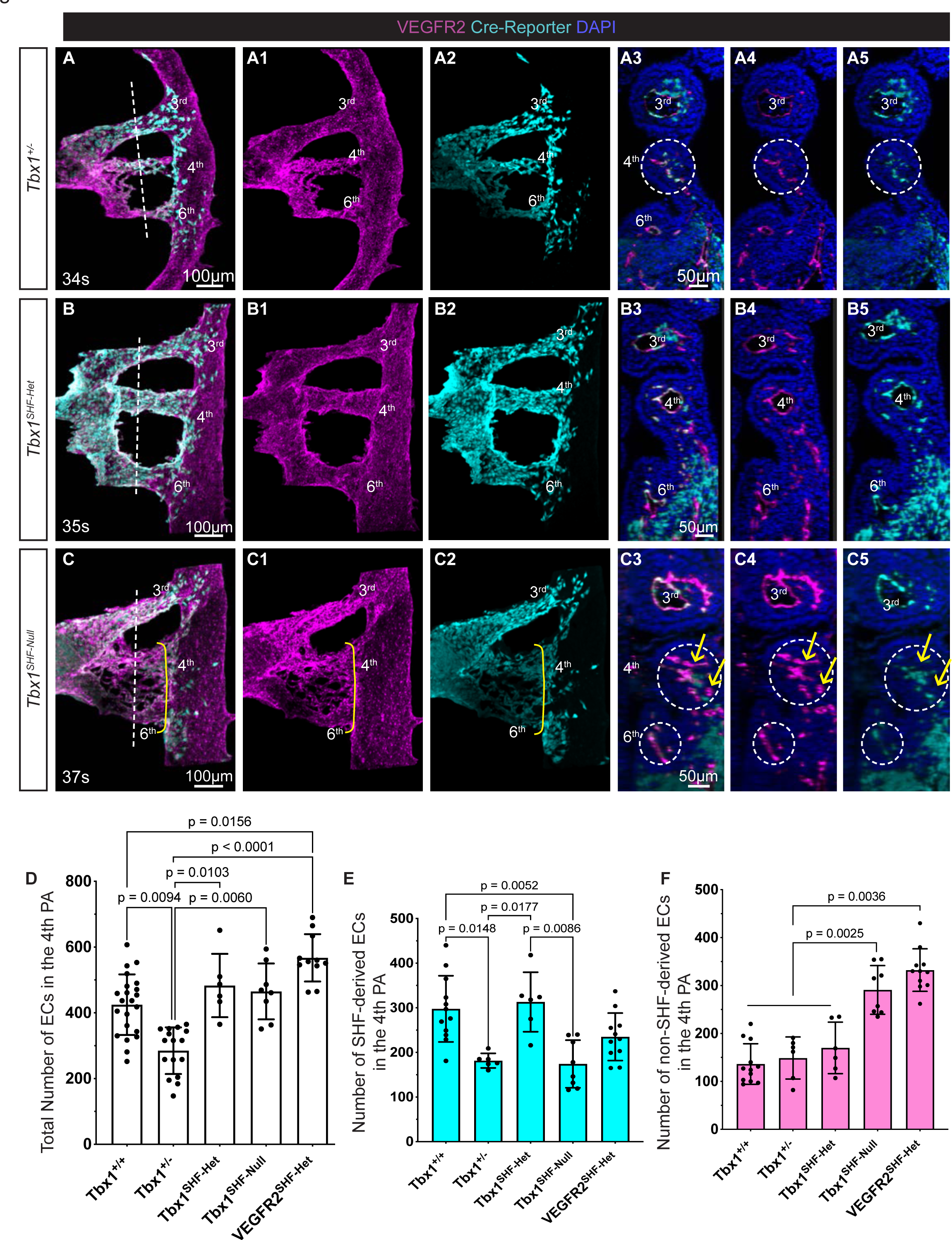
*Tbx1* regulates the number of SHF-derived ECs in 4th pharyngeal arches and EC compensation. Whole mount immunofluorescence staining detecting VEGFR2 (magenta) and Cre-reporter (cyan) in E10.5 embryos. Imaris was used to segment the dorsal aorta and pharyngeal arch ECs and 3D reconstructions: **A-A2**, *Tbx1^+/-^* **B-B2**, *Tbx1^SHF-Het^* and **C-C2**, *Tbx1^SHF-Null^* embryos; The only Cre-reporter+ cells shown are those that lie within the segmented endothelium. Pharyngeal arches are numbered. Yellow brackets mark endothelium in the 4th and 6th arches in **C-C2**. Vertical dashed lines mark the planes of coronal optical sections in **A3-A5, B3-B5, C3-B5**, where all cells are shown. Dashed circles in **A3-A5** and **C3-C5** mark pharyngeal arches. Yellow arrows in **C3-C5** point to VEGFR2+ lineage-negative compensatory ECs. Quantification of total EC numbers, **D**, SHF-derived ECs, **E,** and non-SHF-derived ECs**, F.** Each dot marks one pharyngeal arch, error bars show standard deviations; p values were calculated using 2-tailed, unpaired, non-parametric Kruskal-Wallis tests with Dunn’s correction for multiple testing; only p<0.05 are displayed.

The deletion of a single *Tbx1* allele in the SHF alone (Tbx1^f/+^; Mef2C-AHF-Cre embryos, here referred to as Tbx1^SHF-Het^) did not impact PAA formation, the total number of 4^th^ arch ECs, or the number of SHF-derived ECs in the 4^th^ pharyngeal arch (**Fig. 5B-B5, 5D-E**). This is consistent with earlier studies indicating that the expression of just a single *Tbx1* allele in the mesoderm is sufficient for the development and remodeling of the 4^th^ PAAs ^32^. These data indicate that 1) the heterozygosity of *Tbx1* in the SHF-derived ECs does not impair PAA formation *per se*, and 2) the expression of *Tbx1* in tissues other than the SHF regulates SHF-derived EC number when only one *Tbx1* allele is deleted in the SHF. The latter is consistent with previous studies demonstrating the importance of *Tbx1* expression in the pharyngeal epithelia, the endoderm and the surface ectoderm, in regulating the formation of the 4^th^ PAAs ^32^.

Neither the total EC number nor the number of SHF-derived cells was affected in the 6^th^ pharyngeal arches of *Tbx1^+/-^* embryos (**Fig. S3A-C**), consistent with prior studies demonstrating that the 6^th^ PAAs and the ductus arteriosus, the derivative of the left 6^th^ PAA form in *Tbx1^+/-^* embryos ^24–26^. Together, our data show that *Tbx1* regulates EC number in the 4^th^ pharyngeal arches and that this regulation is non-SHF-cell-autonomous.

Interestingly, the number of SHF-derived ECs in the 4^th^ pharyngeal arch of *Tbx1^+/-^* and *VEGFR2^SHF-Het^* embryos were comparable, and both were diminished to a similar extent compared with their littermate controls (**Fig. 5E**, **Fig. 2I**). But, unlike in *Tbx1^+/-^*embryos, the decrease in the SHF-derived ECs in *VEGFR2^SHF-Het^*embryos was compensated by increased numbers of non-SHF-derived ECs (**Fig. 5F**, 5^th^ bar). In contrast, no such increase in non-SHF-derived ECs was observed in *Tbx1^+/-^* embryos (**Fig. 5F**, 2^nd^ bar). And while only 1 of 11 (9%) *VEGFR2^SHF-Het^* embryos developed a defective 4^th^ PAA at E10.5 and 1 of 21 (4.8%) *VEGFR2^SHF-Het^* embryos developed a 4^th^ PAA-related defect (**Table S6**), 75-100% of *Tbx1^+/-^* embryos have aberrantly formed 4^th^ PAAs at E10.5 – E11.5 ^31,33,34^, and ∼46% of *Tbx1^+/-^* embryos have defective 4^th^ PAA-derived vessels at ≥E13.5, the time when PAA remodeling is complete ^31,33,34^.

Milder PAA-related phenotypes in *VEGFR2^SHF-Het^* embryos compared with the *Tbx1^+/-^* correlate with the presence of compensatory ECs in the *VEGFR2^SHF-Het^* model, suggesting that the compensatory endothelium can rescue PAA formation and remodeling. To determine the cell type(s) in which *Tbx1* was required to elicit the compensatory endothelium, we generated Tbx1^f/f^; Mef2C-AHF-Cre embryos (*Tbx1^SHF-Null^*). The numbers of SHF-derived cells in *Tbx1^+/-^ and Tbx1^SHF-Null^* embryos were comparable, and both were significantly decreased (by ∼40%) compared to *Tbx1^+/+^* controls (**Figs. 5E**); However, unlike in *Tbx1^+/-^* embryos, the total number of 4^th^ pharyngeal arch ECs in *Tbx1^SHF-Null^* embryos was complemented by the recruitment of the compensatory endothelium (**Fig. 5D-F**).

To get insights into the cellular origin of the compensatory endothelium in *Tbx1^SHF-Null^* embryos, we examined the 3D connectivity of vessels surrounding the pharyngeal arches (**Fig. 6**). Unlike *VEGFR2^SHF-Het^* and *VEGFR2^SHF-KO^*embryos, *Tbx1^+/-^* embryos lacked compensatory ECs, and while prominent sprouts can be seen in *VEGFR2^SHF-KO^* embryos (arrows in **Fig 6B1-B4**), sprouts extending from veins into the pharyngeal arches were absent in *Tbx1^+/-^* embryos: compare serial optical sections in **Fig. 6A-A5** with **Fig. 6C-C4**, noting smooth CV surface marked with red asterisks. In contrast, conspicuous venous sprouts extending from the cardinal and pharyngeal veins into the 4^th^ pharyngeal arches were observed in *Tbx1^SHF-Null^* embryos that, like *VEGFR2^SHF-Het^* and *VEGFR2^SHF-KO^* embryos, also contained compensatory ECs (**Fig. 6E-E4**, arrows in **6E2-E3**). Our experiments collectively demonstrate that *Tbx1* expression in both the SHF and non-SHF lineages governs the abundance of SHF-derived ECs. Notably, *Tbx1* also influences the recruitment of compensatory endothelium, doing so in a non-SHF-cell-autonomous manner. Our studies also suggest that the compensatory ECs arise via angiogenic sprouting from veins and that wild-type levels of *Tbx1* in tissues other than the SHF-derived mesoderm regulate this compensatory response.

**Figure 6.**
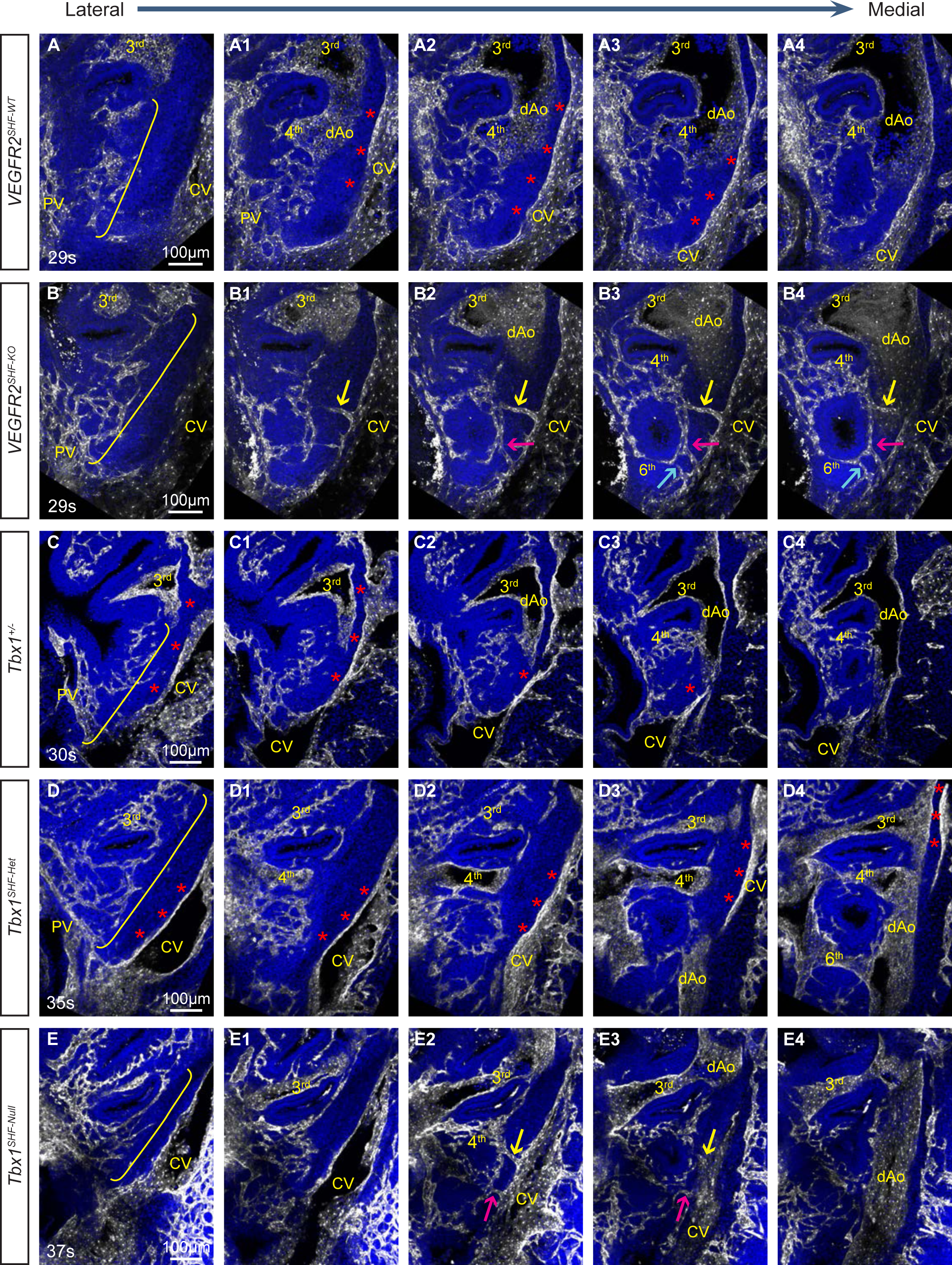
Ectopic sprouts from veins extend into pharyngeal arches of *VEGFR2^SHF-KO^* and *Tbx1^SHF-Null^* but not *Tbx1^+/-^* embryos. A-C, E10.0 and **D-E,** E10.5 whole embryos stained to detect VEGFR2 (white) and nuclei (blue); 20μm-thick serial sagittal optical sections are shown in the lateral-to-medial order. Yellow brackets in **A-E** mark pharyngeal endothelial plexus. Arches are numbered. Red asterisks mark smooth surfaces of cardinal veins (CV). Arrows of the same color follow the same sprout in different serial sections. dAo - dorsal aorta, PV - pharyngeal vein.

### Angiogenic sprouting from veins gives rise to the compensatory endothelium

The presence of ectopic sprouts extending from the CV and PV into the pharyngeal arches of *VEGFR2^SHF-Het^* and *VEGFR2^SHF-KO^* embryos suggested that compensatory endothelium is derived from veins. To test this hypothesis, we used lineage tracing to label venous endothelium in *VEGFR2^SHF-WT^, VEGFR2^SHF-Het^,* and *VEGFR2^SHF-KO^* embryos. To check the specificity of venous labeling, we first employed the *Apj-CreERT2* transgenic strain ^35^ and tamoxifen injection at E9.5 to induce Cre-mediated recombination in veins. Embryos were dissected at E10.5, stained, and imaged to detect the entire vasculature in and around the pharyngeal region (**Fig. S3D-E**). We observed extensive labeling of venous vasculature, including the PV, CV, and venous intersomitic branches in wild-type embryos (**Fig. S3D**). In contrast, arterial structures, including the dorsal aorta, arterial intersomitic branches, and PAAs, were not labeled except for one or two cells (**Fig. S3E**). These data indicated high specificity for venous lineage labeling using the *Apj-CreERT2* transgenic strain when tamoxifen was injected at E9.5. Thus, Apj-directed gene expression is highly specific to veins at E9.5.

To label venous vasculature in embryos with the conditional, Mef2C-AHF-Cre-mediated inactivation of *VEGFR2* in the SHF, we generated the *Apj^DreERT^*^2^*^/+^*knock-in mice and crossed them to obtain *VEGFR2^SHF-WT^; Apj^DreERT^*^2^*^/+^*, *VEGFR2^SHF-Het^; Apj^DreERT^*^2^*^/+^*, and *VEGFR2^SHF-KO^; Apj^DreERT^*^2^*^/+^*embryos carrying a fluorescent Dre-reporter. This approach allows conditional labeling of venous cells independently of modifying VEGFR2 expression in the SHF using the Mef2C-AHF-Cre transgene. Following tamoxifen injection at E9.5, we dissected embryos at E10.5 and analyzed the contribution of Apj-lineage-derived cells to the vasculature in controls and mutants (**Fig. 7**). We found that lineage labeling efficiency using a single injection of tamoxifen in the *Apj^DreERT^*^2^*^/+^* knock-in strain was ∼15% across all genotypic classes (**Fig. S3F-G**). The specificity of venous blood vessel labeling was high since we did not see lineage-labeled cells in the dorsal aorta or its branches (**Fig. 7A-B**).

**Figure 7.**
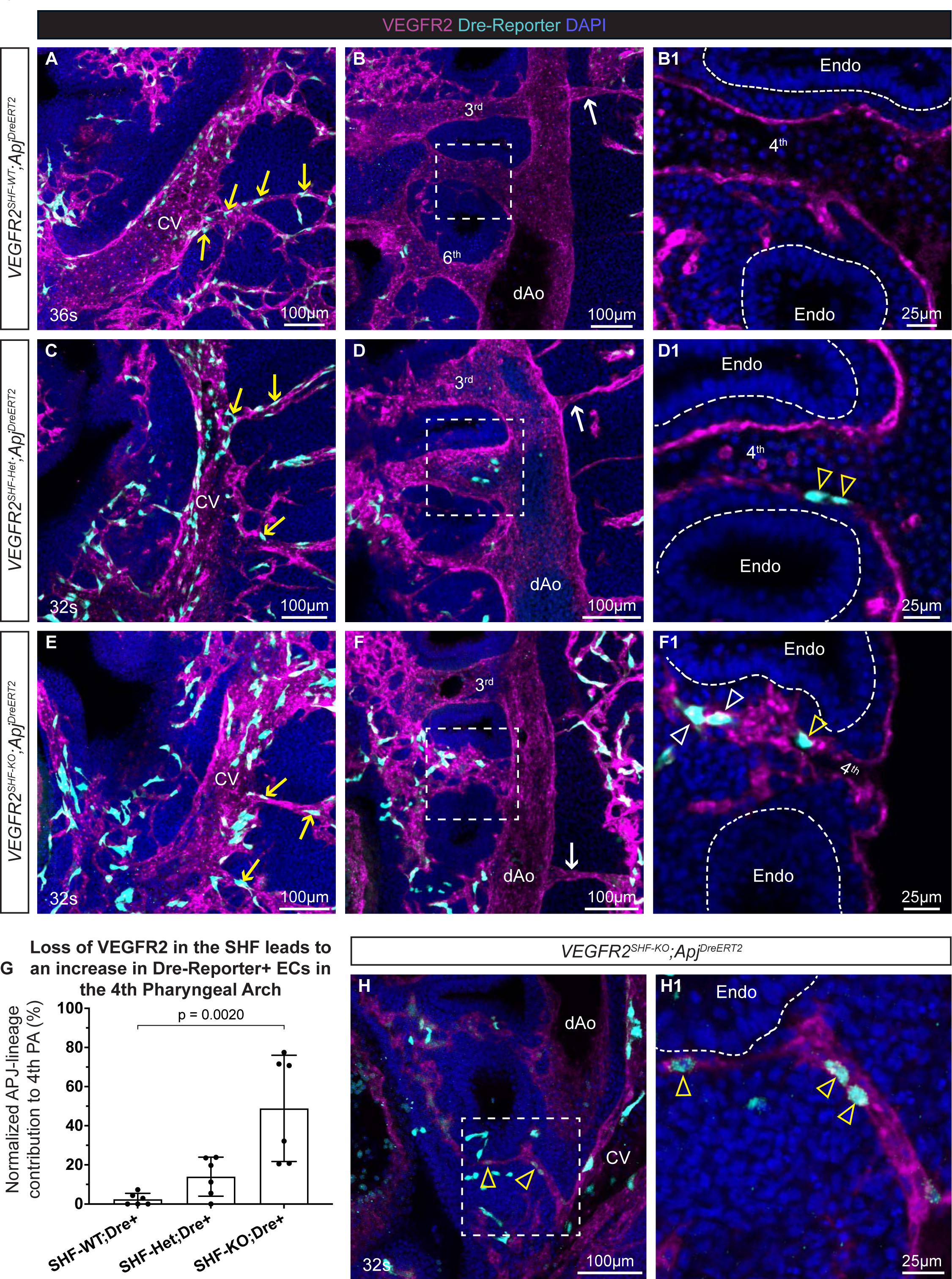
Angiogenic sprouting from veins gives rise to the compensatory endothelium. Dre-rox recombination was used to fate map cells of *Apj*-lineage in *VEGFR2^SHF-WT^*, *VEGFR2^SHF-Het^,* and *VEGFR2^SHF-KO^* embryos. Tamoxifen was injected at E9.5, and embryos were dissected at E10.5. **A-F,** Whole mount immunofluorescence staining detecting Dre-reporter (cyan) and VEGFR2 (magenta). 50μm-thick sagittal optical sections show *Apj^DreERT^*^2^ lineage labeling in the cardinal vein (CV) and intersomitic veins (yellow arrows) in **A, C,** and **E** but not the in the dorsal aorta (dAo) or intersomitic arteries (white arrows) in **B, D,** and **F** of *VEGFR2^SHF-WT^*, *VEGFR2^SHF-Het^,* and *VEGFR2^SHF-KO^*embryos. Boxes in **B, D** and **F** are expanded to the right to show *Apj^DreERT^*^2^ lineage-labeled ECs in thin, 0.45 μm optical sections. Note Apj-lineage cells lining the lumen of the mature 4th PAA in *VEGFR2^SHF-Het^* embryo, arrowheads in **D1**, and the lumen of the forming 4th PAA in *VEGFR2^SHF-KO^* embryo, yellow arrowhead in **F1.** White arrowheads point to Apj-lineage cells in the 4th pharyngeal arch plexus, **F1**. **G.** Total number of Apj-lineage cells per 4th arch. Each dot marks one arch. p values were calculated using 2-tailed, unpaired, non-parametric Kruskal-Wallis tests with Dunn’s correction for multiple testing; only p<0.05 are displayed. **H.** 50μm-thick sagittal optical section showing CV sprout extending into the pharyngeal arch region. Boxed region in **H** is expanded in **H1** as a 2 μm thick optical section. Arrowheads point to Apj-lineage cells (cyan). Endoderm (endo) layers are marked by dotted lines.

Consistent with the observation of ectopic venous sprouts and the presence of venous marker-expressing cells in the pharyngeal arches of *VEGFR2^SHF-Het^*and *VEGFR2^SHF-KO^* mutants, we observed increased numbers of Apj-lineage labeled cells in the pharyngeal arch endothelium in *VEGFR2^SHF-Het^; Apj^DreERT^*^2^*^/+^* and *VEGFR2^SHF-KO^; Apj^DreERT^*^2^*^/+^* embryos compared with controls (compare **Fig. 7D-D1** and **7F-F1** with **7B-B1**). Quantification of all Apj-lineage labeling of ECs in the 4^th^ pharyngeal arches revealed a trend and a significant increase in *Apj*-Dre-labeled ECs in *VEGFR2^SHF-Het^* and *VEGFR2^SHF-KO^* embryos, respectively, compared with *VEGFR2^SHF-WT^*embryos (**Fig. 7G**). In *VEGFR2^SHF-Het^* embryos, Apj-lineage cells were observed in the mature lumen of the 4^th^ PAA (**Fig. 7D-D1**, arrowheads). In *VEGFR2^SHF-KO^* embryos, Apj-lineage cells were seen both in the plexus endothelium (**Fig. 7F1**, white arrowheads) as well as in the wall of the thin but lumenized portion of the forming 4^th^ PAA (**Fig. 7F1**, yellow arrowhead). Sprouts emanating from the cardinal vein also contained Apj-lineage labeled cells (arrowheads in **Fig. 7H-H1**). Together, these data show that cells of venous lineage give rise to the compensatory endothelium in *VEGFR2^SHF-Het^* and *VEGFR2^SHF-KO^* embryos.

### *VEGF-A* is upregulated in the pharyngeal arches of *VEGFR2^SHF-KO^* embryos but not *Tbx1^+/-^* embryos

The experiments described above show that compensatory ECs in the 4th pharyngeal arches of *VEGFR2^SHF-Het^*and *VEGFR2^SHF-KO^* mutants arise from veins. We hypothesized that the reduction/loss of SHF-derived ECs in the pharyngeal arches of these mutants could induce the expression of proangiogenic factors, such as VEGF-A, eliciting angiogenic response in the nearby veins. To test this idea, we performed single-molecule (sm) RNA-FISH to examine *VEGF-A* mRNA levels at early E9.5 (23-24s) before compensatory ECs extensively colonized pharyngeal arches. Quantification showed that *VEGF-A* mRNA expression was significantly increased in the SHF and trended upwards in the endoderm of *VEGFR2^SHF-KO^*embryos compared to *VEGFR2^SHF-WT^* (**Fig. S4A-A1, B-B1,** quantified in **S4E-F**). In contrast, *VEGF-A* mRNA was not upregulated in *Tbx1*^+/-^ embryos (**Fig. S4C-C1, 4D-D1,** quantified in **S4G-H**). These data suggest that *VEGF-A* contributes to eliciting the angiogenic response from the nearby veins and that *Tbx1* potentially regulates its expression.

### VEGFR2 expression in veins is required for the compensatory response

VEGF-A regulates angiogenesis by activating VEGFR2 signaling ^36^. We hypothesized that the inactivation of VEGFR2 in veins would block angiogenic response and that the deletion of VEGFR2 both in the SHF and the veins would block pharyngeal arch vascularization. To test these hypotheses, we employed the Apj-CreERT2 transgenic line. Although tamoxifen injection at E9.5 in this strain results in specific labeling of veins (**Fig. 7A**, **Fig. S3D**), injection at E8.5 labels both the vein and SHF (**Fig. S5**). This is consistent with the expression pattern of *Apj* mRNA ^37,38^. Indeed, when tamoxifen was injected at E8.5, both the cardinal vein and SHF expressed *Rosa^tdTomato^* Cre-reporter at E10.5 (**Fig. S5**).

To determine whether expression of VEGFR2 in veins was required for the presence of compensatory ECs in the 4^th^ pharyngeal arches, we crossed *VEGFR2^+/-^; Apj-CreERT2* transgenic males with *VEGFR2^flox/flox^*; *Rosa^tdTom/tdTom^* females to generate *VEGFR2^flox/-^; Apj-CreERT2+; Rosa^tdTom/+^* (*VEGFR2^Apj-KO^*) embryos. *VEGFR2^+/-^; Apj-CreERT2* males were also crossed with *Rosa^tdTom/tdTom^*females to generate *VEGFR2^+/+^; Apj-CreERT2 +; Rosa^tdTom/+^*(*VEGFR2^Apj-WT^*) controls. *VEGFR2^Apj-KO^* embryos degenerated between E10.5 and E10.75. Therefore, we dissected embryos earlier, in the morning of E10.5, at the 28-29 somite stage, to analyze the vasculature in live embryos. The 28-29 somite stage is suitable for investigating compensatory EC response because compensatory ECs are readily observed in *VEGFR2^SHF-KO^* mutants at this stage (**Fig. 4E-E1, H-H1**).

To check the ablation of VEGFR2 in the Apj lineage following the injection of tamoxifen at E8.5, we assayed the expression of VEGFR2 in veins and the SHF in *VEGFR2^Apj-WT^*and *VEGFR2^Apj-KO^* embryos (**Fig. S5**). In *VEGFR2^Apj-WT^* embryos, Cre-reporter+ cells in the anterior cardinal vein and SHF expressed VEGFR2 as expected (**Fig. S5A-A3, S5C-C2**). In *VEGFR2^Apj-KO^* mutants, VEGFR2 was downregulated in the anterior cardinal vein and SHF (**Fig. S5B-B3, S5D-D2,** arrowheads), although some escapee Cre-reporter+ venous ECs expressing VEGFR2 were also present (**Fig. S5B1-B3**, arrows).

Most of the 4^th^ pharyngeal arch endothelium arose from the Apj-lineage in *VEGFR2^Apj-WT^* controls (**Fig. 8A-A3**, quantified in **8C**). Remarkably, it was mainly the escapee cells still expressing VEGFR2 in the Apj lineage that were able to contribute to the pharyngeal arch endothelium in *VEGFR2^Apj-KO^* mutants (**Fig. 8B-B3**, yellow arrows; quantified in **8C**). This observation is consistent with earlier studies demonstrating that the absence of VEGFR2 precluded the contribution of progenitors to embryonic endothelium ^39,40^, and with studies in this manuscript demonstrating that the expression of VEGFR2 in the SHF was stringently required for the contribution of SHF-derived cells to the pharyngeal arch artery endothelium (**Fig. 2**).

**Figure 8.**
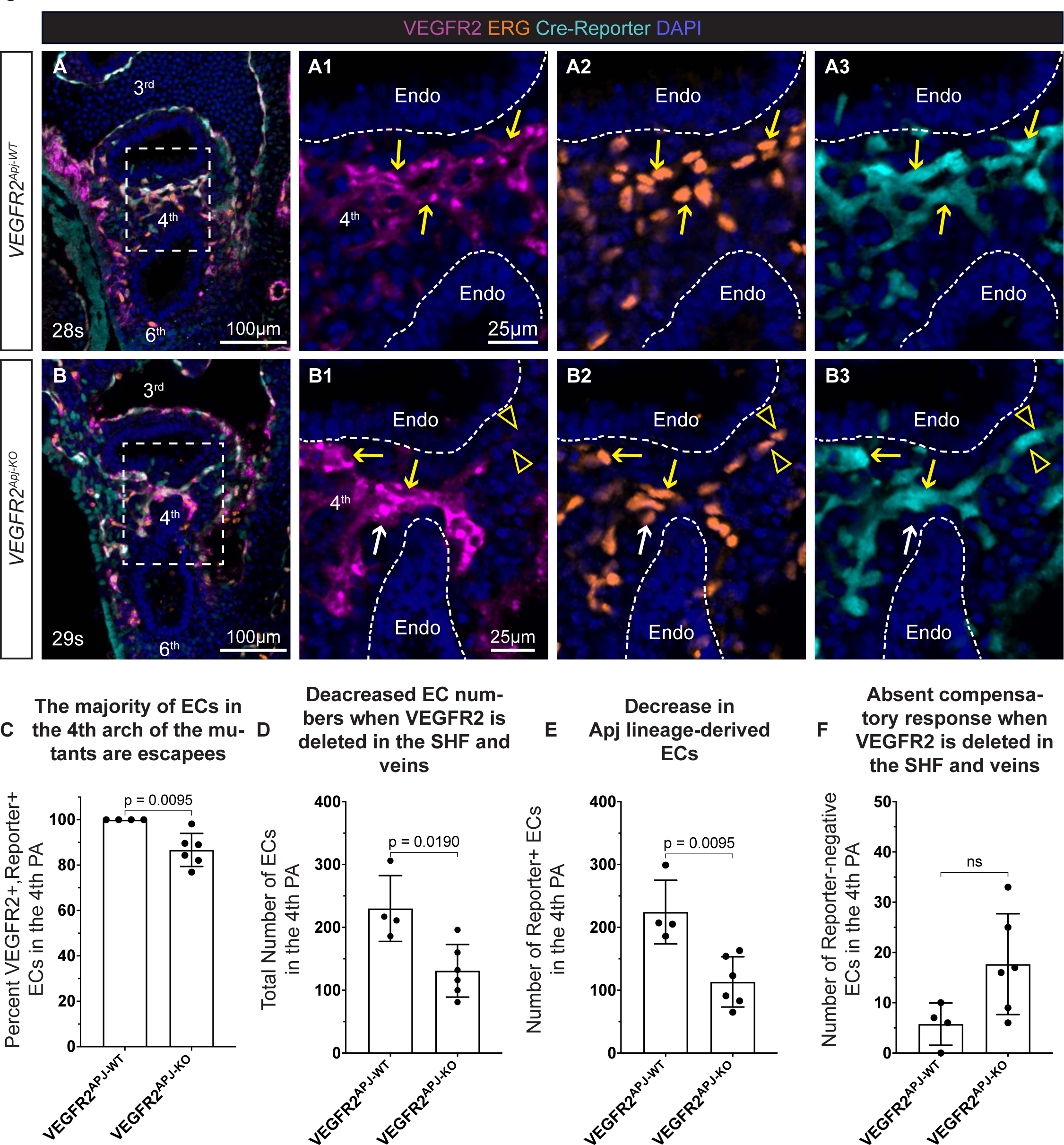
The deletion of VEGFR2 in the veins significantly attenuates compensatory response. VEGFR2 was ablated in the SHF and veins in VEGFR2^+/+^; Apj-CreERT2^+^; Rosa^tdTom/+^ (VEGFR2^Apj-WT^) and VEGFR2^flox/-^; Apj-CreERT2^+^; Rosa^tdTom/+^ (VEGFR2^Apj-KO^) embryos by injecting tamoxifen at E8.5. For the demonstration of efficient deletion of VEGFR2 the SHF and veins, see **Fig. S5**. Whole embryos were stained by immunofluorescence and imaged by confocal microscopy. **A-A3,** 0.45 μm sagittal optical sections from VEGFR2^Apj-WT^ and **B-B3,** VEGFR2^Apj-KO^ embryos. Boxed areas in **A** and **B** are expanded in **A1-A3** and **B1-B3**. Yellow arrows point to VEGFR2+ Apj-lineage cells (escapees); arrowheads point to the rare VEGFR2-negative Apj-lineage cells, and white arrow points to a non-Apj-lineage VEGFR2+ cell). Endoderm (endo) is bordering the 4th arch is outlined. **C-F**, Quantification of ECs in the 4th pharyngeal arches. **C,** The majority of 4th pharyngeal arch ECs are escapees expressing VEGFR2 in VEGFR2^Apj-KO^ mutants. **D,** Significant decrease in the total number of ECs in VEGFR2^Apj-KO^ embryos. **E,** Significant decrease in the Apj-lineage ECs in the 4th arch of VEGFR2^Apj-KO^ mutants. **F,** A meager compensatory response in the 4th arches of VEGFR2^Apj-KO^ mutants does not bring the total number of ECs to control levels. Each dot marks one pharyngeal arch, all data points are shown, as well as means and standard deviations; p values were calculated using 2-tailed, unpaired, non-parametric Mann-Whitney tests.

Overall, there was ∼ a two-fold decrease both in the total number of ECs (**Fig. 8D**) and in the number of Apj-derived ECs (**Fig. 8E**) in *VEGFR2^Apj-KO^*mutants compared with *VEGFR2^Apj-WT^* controls. Although some compensatory ECs were present in *VEGFR2^Apj-KO^* mutants (**Fig. 8B1-B3**, white arrow), they numbered in dozens rather than hundreds, as in *VEGFR2^SHF-Het^* embryos (compare **Fig. 8F** and **Fig. 5F**, last bar). This compensatory response was not sufficient to reconstitute the number of pharyngeal arch ECs in *VEGFR2^Apj-KO^* mutants to the levels seen in *VEGFR2^Apj-WT^* controls. Thus, the deletion of VEGFR2 in veins effectively inhibited the compensatory endothelial response, resulting in EC deficiency in the pharyngeal arches followed by embryonic demise.

## Discussion

Proper development and remodeling of the PAAs, particularly the left 4^th^ PAA, is crucial for establishing correct blood circulation after birth. For reasons that are not well understood, the 4^th^ PAA pair is particularly sensitive to genetic mutations and environmental insults. In this manuscript, we describe the discovery of an embryonic resiliency mechanism facilitating the recovery of the 4^th^ PAAs and their derivatives when EC progenitors of the PAAs are depleted.

The constancy of development, i.e., the development of stereotypical organs and body plans despite their incredible complexity, has captivated and confounded researchers. In 1942, Waddington proposed the concept of buffering to explain this remarkable characteristic of multicellular organisms. He hypothesized that there exists a buffering capacity based on an optimum and a threshold stimuli, beyond which the optimal response can no longer be achieved ^1^. However, despite a few notable studies ^41–43^, our understanding of buffering processes and their underlying mechanisms is far from complete.

By carefully quantifying pharyngeal arch EC numbers, we observed that the average number of total ECs in the 4^th^ pharyngeal arches was similar across multiple different studies utilizing control mice and genotypes examined in our lab by independent investigators over the past decade (∼476 ± 81 ECs per 4^th^ arch) ^8^. We also noticed that a decrease in this number by 25-33% in three different mutant models was associated with defective formation and remodeling of the 4^th^ PAAs. The models we examined were *Tbx1^+/-^* mice (this manuscript) and mice in which either fibronectin (Fn1) or integrin α5 were downregulated in the Isl1 lineage ^8,44^. It is unlikely that the 4^th^ PAA defects in these models arose due to the roles of these proteins in PAA formation *per se*. This is because the expression of integrin α5 in PAA in progenitors or in the endothelium is not required for PAA formation ^45,46^. Similarly, the deletion of one *Tbx1* allele in PAA progenitors or the endothelium does not interfere with PAA formation and remodeling ^32^. Since neither the expression of integrin α5 nor the presence of two copies of the *Tbx1* gene is required for the endothelium to form the 4^th^ PAAs, our data suggest that the presence of the proper number of ECs in the pharyngeal arches is important. Thus, we propose that there exists an optimum total number of ECs in the 4^th^ pharyngeal arches, and a 25-33% decrease in this number leads to unpredictable developmental trajectories of PAAs and their derivatives, resulting in lethal CHD such as IAA-B.

Prior studies showed that the expression of VEGFR2 is required for progenitors to adopt EC fate ^39,40,47,48^. Therefore, we deleted VEGFR2 in the SHF as a means to ablate SHF-derived ECs. This strategy worked well; we achieved an efficient deletion of VEGFR2 in the SHF in *VEGFR2^SHF-KO^* embryos, and the contribution of SHF-derived cells to pharyngeal arch ECs was severely impaired. These experiments allowed us to discover the existence of a compensatory mechanism that can rescue PAA formation and remodeling.

Despite similar total numbers of ECs in the pharyngeal arches in *VEGFR2^SHF-Het^*and *VEGFR2^SHF-KO^* embryos, *VEGFR2^SHF-KO^* embryos displayed more severe defects in PAA formation and remodeling. A notable difference between these genotypic classes is a near-complete absence of SHF-derived ECs in the 4^th^ pharyngeal arches of *VEGFR2^SHF-KO^* embryos. These findings highlight the essential role of the SHF as the main source of 4^th^ PAA ECs and show that in addition to the compensatory endothelium, a critical number of SHF-derived ECs is necessary for the proper formation and remodeling of the fourth PAAs. This could be due to a delayed accrual of pharyngeal arch ECs in *VEGFR2^SHF-KO^* embryos compared with *VEGFR2^SHF-WT^* and *VEGFR2^SHF-Het^* and delayed reprogramming of venous cells into arterial. Together, our data suggest the importance of maintaining a specific threshold of SHF-derived ECs for the constancy of PAA development and remodeling.

Our studies suggest that crosstalk between SHF-derived ECs and the surrounding pharyngeal tissues regulates the addition of compensating endothelium to the PAAs. A crosstalk between the endothelium and the developing tissues has been implicated in organ development before ^49–51^. Thus, we hypothesize that the presence of an optimal number of SHF-derived ECs suppresses ectopic angiogenesis from veins, resulting in the formation of SHF-derived PAAs. This is because even in control embryos, *VEGF-A* mRNA is highly expressed in the endoderm and the SHF, yet no ectopic venous sprouts extending into the pharyngeal arches are observed when the proper number of SHF-derived ECs is present. However, when the number of SHF-derived ECs is lowered, the pharyngeal tissue microenvironment is modified in some way, resulting in increased synthesis of *VEGF-A* and likely additional other factors, and maybe in the downregulation of angiogenesis inhibitors, altogether inducing an ectopic angiogenic response from the nearby veins.

We don’t know whether SHF-derived and vein-derived ECs are equivalent in their function within adult PAA-derived vessels. However, there are multiple examples in the literature where veins give rise to arteries in mice and fish during the physiological process of development. In mice, coronary arteries on the dorsal side of the heart are derived from reprogrammed venous cells arising via angiogenesis from the sinus venosus ^52,53^. Additional physiological examples of veins giving rise to arteries include retinal angiogenesis and early mouse development ^54–57^. In zebrafish, live imaging and lineage tracing showed that angiogenesis from veins gives rise to basilar, central, and nasal ciliary arteries in the brain ^58,59^. Angiogenic sprouting from veins also contributes to the growing arteries in pectoral and caudal fins during zebrafish development and regeneration ^55,60–62^. Thus, the ablation of SHF-derived endothelium in the pharyngeal arches may have evoked an inherent capacity of veins to sprout and provide the endothelium to rescue arch artery development.

We hypothesize that there may be an evolutionary advantage for the formation of the PAAs from SHF-derived progenitors rather than veins. Possibly, this could allow for an efficient coordination of cardiac and vascular development and proper routing of the blood. In addition to the PAA endothelium, SHF gives rise to the endocardium and myocardium of the right ventricle and the cardiac outflow tract ^47,63–67^. The PAA-derived vessels are contiguous with the cardiac outflow tract-derived vasculature, the ascending aorta, and the pulmonary trunk. Thus, the proper PAA connectivity with the cardiac structures may necessitate common regulatory mechanisms (e.g., *Tbx1*, Mef2C-AHF enhancer) and progenitor sources (e.g., the SHF). Finally, PAAs have to form and connect the heart with the dorsal aorta in a timely manner to ensure that PAA-derived vessels, the aortic arch, and the ductus are correctly connected with the heart. SHF-derived EC progenitors in the lateral plate mesoderm are incorporated into the pharyngeal arches, providing PAA EC source as early as E9.0, while the recruitment of ECs via angiogenesis from veins requires the time for these cells to migrate into the pharyngeal arches, reprogram to the arterial cell fate, and to form lumenized arterial vessels that connect to the arterial outflow vasculature and the dorsal aortae. This connectivity is essential for routing blood into the systemic circulation and a delay in PAA formation may undermine the timely developing and remodeling of this complex vascular circuit.

Most patients with 4^th^ PAA-derived CHD, including IAA-B, are diagnosed with 22q11DS ^20,22^. Studies using *Tbx1^+/-^* mice as a model system demonstrated that defects in the formation of the 4^th^ PAAs are attributed to haploinsufficiency in the *Tbx1* gene ^24–26,32,68^; however, mechanisms by which *Tbx1* regulates PAA formation have remained unclear. Experiments in this manuscript show that *Tbx1* is a critical regulator of SHF-derived EC numbers and the compensatory response from veins.

Consistent with prior reports ^31^, we found that the 4^th^ PAAs in *Tbx1^+/-^* embryos were either hypoplastic or aplastic. Despite the high penetrance of 4^th^ PAA formation defects in *Tbx1^+/-^* embryos (75-100%), about half of *Tbx1^+/-^*embryos are viable and fertile ^31,33,34^. The partial recovery of 4^th^ PAA defects is dependent on whether the 4^th^ PAAs are aplastic (meaning only plexus ECs are present at E10.5/E11.5 in the 4^th^ arches) or hypoplastic (a thin PAA connecting with the ventral and the dorsal aortae is present) in *Tbx1^+/-^* embryos ^31^. The proportion of *Tbx1^+/-^*embryos surviving to adulthood without 4^th^ PAA-related defects is equal to the percentage of embryos with hypoplastic 4^th^ PAAs ^31^. In other words, thin 4^th^ PAAs in *Tbx1^+/-^*embryos can recover and give rise to their derivative adult vessels. But if the 4^th^ arch endothelium remained in the form of plexus by E11.5 (e.g., a collection of tiny capillaries without the presence of a central lumenized vessel in the 4th arch), this resulted in 4^th^ PAA-related defects such as IAA-B or RERSA.

Although the reason for the variability of whether plexus ECs stay as a collection of capillaries rather than morphing into a central albeit thin arch artery in *Tbx1^+/-^* embryos is unknown, it could be related to a potential variability in the numbers of SHF-derived ECs or variability in the compensatory response among different *Tbx1^+/-^* embryos. The PAA recovery mechanisms in *Tbx1^+/-^* embryos and the regulation of compensatory endothelium by *Tbx1* need to be more thoroughly investigated to answer these questions.

Taken together, our studies revealed a novel mechanism of embryonic resiliency and vascular plasticity and highlighted the importance of vein-derived compensatory endothelium for PAA development and neonatal survival.

## Supporting information

Supplemental statistical analysis

## MATERIALS AND METHODS

All Materials and Methods are detailed in the Supplemental Methods file.

## Author contributions

AJR and SA conceived and designed the experiments; AJR, CAV, and HZ performed the experiments; KE performed statistical analyses of proportions, KD processed embryos for micro-CT imaging and helped with the analyses and interpretation of congenital heart defects. AJR and SA analyzed the data and wrote the paper.

## Acknowledgments

We thank Cecilia Arriagada for the graphical abstract. Caolán O’Donnell and members of the Astrof lab for helpful suggestions throughout this work. Brittany Ramirez for her artistic expertise. This work was supported by funding from the National Heart, Lung, and Blood Institute of the National Institutes of Health grants R01 HL103920, R01 HL134935, and R01 HL158049, and a Transformational Project Award from the American Heart Association 20TPA35490074 to SA. AJR was supported by the National Heart, Lung, and Blood Institute of the National Institutes of Health pre-doctoral fellowships F31 HL150949, HL103920-08S1, and by the National Institute of Arthritis and Musculoskeletal and Skin Diseases training grant T32052273-11. CAV was partly supported by the NJMS M.D./Ph.D. program at Rutgers University. HZ was supported by the AHA Postdoctoral Fellowship 23POST1022380.

## References

1. Waddington CH. Canalization of development and the inheritance of the acquired characters. Nature. 1942;150:563–564.

2. High FA, Jain R, Stoller JZ, Antonucci NB, Lu MM, Loomes KM, Kaestner KH, Pear WS, Epstein JA. Murine Jagged1/Notch signaling in the second heart field orchestrates Fgf8 expression and tissue-tissue interactions during outflow tract development. J Clin Invest. 2009;119:1986–1996. doi: 10.1172/JCI38922

3. Stoller JZ, Epstein JA. Cardiac neural crest. Semin Cell Dev Biol. 2005;16:704–715. doi: 10.1016/j.semcdb.2005.06.004

4. Benjamin EJ, Virani SS, Callaway CW, Chamberlain AM, Chang AR, Cheng S, Chiuve SE, Cushman M, Delling FN, Deo R, et al. Heart Disease and Stroke Statistics—2018 Update: A Report From the American Heart Association. In: *Circulation*. 2018:e67-e492.

5. Tsao CW, Aday AW, Almarzooq ZI, Alonso A, Beaton AZ, Bittencourt MS, Boehme AK, Buxton AE, Carson AP, Commodore-Mensah Y, et al. Heart Disease and Stroke Statistics-2022 Update: A Report From the American Heart Association. Circulation. 2022:CIR0000000000001052. doi: 10.1161/CIR.0000000000001052

6. Graham A. Development of the pharyngeal arches. Am J Med Genet A. 2003;119A:251–256. doi: 10.1002/ajmg.a.10980

7. Wang X, Chen D, Chen K, Jubran A, Ramirez A, Astrof S. Endothelium in the pharyngeal arches 3, 4 and 6 is derived from the second heart field. Dev Biol. 2017;421:108–117. doi: 10.1016/j.ydbio.2016.12.010

8. Warkala M, Chen D, Ramirez A, Jubran A, Schonning M, Wang X, Zhao H, Astrof S. Cell-Extracellular Matrix Interactions Play Multiple Essential Roles in Aortic Arch Development. Circ Res. 2021;128:e27–e44. doi: 10.1161/CIRCRESAHA.120.318200

9. Nagelberg D, Wang J, Su R, Torres-Vázquez J, Targoff KL, Poss KD, Knaut H. Origin, specification, and plasticity of the great vessels of the heart. In: Current Biology. 2015:2099–2110.

10. Paffett-Lugassy N, Singh R, Nevis KR, Guner-Ataman B, O’Loughlin E, Jahangiri L, Harvey RP, Burns CG, Burns CE. Heart field origin of great vessel precursors relies on nkx2.5-mediated vasculogenesis. In: Nature Cell Biology. 2013:1362–1369.

11. Li P, Pashmforoush M, Sucov HM. Mesodermal retinoic acid signaling regulates endothelial cell coalescence in caudal pharyngeal arch artery vasculogenesis. Dev Biol. 2012;361:116–124. doi: 10.1016/j.ydbio.2011.10.018

12. Kau T, Sinzig M, Gasser J, Lesnik G, Rabitsch E, Celedin S, Eicher W, Illiasch H, Hausegger K. Aortic Development and Anomalies. In: Seminars in Interventional Radiology. 2007:141–152.

13. Celoria G, Patton R. Congenital absence of the aortic arch. American Heart Journal. 1959;58:407–413. doi: 10.1016/0002-8703(59)90157-7

14. Levitt B, Richter JE. Dysphagia lusoria: a comprehensive review. Dis Esophagus. 2007;20:455–460. doi: 10.1111/j.1442-2050.2007.00787.x

15. Kirby ML. Pulmonary atresia or persistent truncus arteriosus: is it important to make the distinction and how do we do it? Circ Res. 2008;103:337–339. doi: 10.1161/CIRCRESAHA.108.174862

16. Ward C, Stadt H, Hutson M, Kirby ML. Ablation of the secondary heart field leads to tetralogy of Fallot and pulmonary atresia. Dev Biol. 2005;284:72–83. doi: 10.1016/j.ydbio.2005.05.003

17. McDonald-McGinn DM, Sullivan KE. Chromosome 22q11.2 deletion syndrome (DiGeorge syndrome/velocardiofacial syndrome). Medicine (Baltimore). 2011;90:1–18. doi: 10.1097/MD.0b013e3182060469

18. McDonald-McGinn DM, Sullivan KE, Marino B, Philip N, Swillen A, Vorstman JA, Zackai EH, Emanuel BS, Vermeesch JR, Morrow BE, et al. 22q11.2 deletion syndrome. Nat Rev Dis Primers. 2015;1:15071. doi: 10.1038/nrdp.2015.71

19. Momma K. Cardiovascular anomalies associated with chromosome 22q11.2 deletion syndrome. Am J Cardiol. 2010;105:1617–1624. doi: 10.1016/j.amjcard.2010.01.333

20. Moerman P, Dumoulin M, Lauweryns J, Van der Hauwaert LG. Interrupted right aortic arch in DiGeorge syndrome. In: Heart. 1987:274–278.

21. Goldmuntz E, Driscol DA, Budarf ML, Zackai E, McDonald-McGinn DM, Biegel JA, Emanuel BS. Microdeletions of chromosomal region 22q11 in patients with congenital conotruncal cardiac defects. Journal of medical genetics. 1993;30:807–812.

22. Van Mierop LHS, Kutsche LM. Cardiovascular anomalies in DiGeorge syndrome and importance of neural crest as a possible pathogenetic factor. The American Journal of Cardiology. 1986;58:133–137. doi: 10.1016/0002-9149(86)90256-0

23. Collins-Nakai RL, Dick M, Parisi-Buckley L, Fyler DC, Castaneda AR. Interrupted aortic arch in infancy. The Journal of Pediatrics. 1976;88:959–962. doi: 10.1016/S0022-3476(76)81049-9

24. Jerome LA, Papaioannou VE. DiGeorge syndrome phenotype in mice mutant for the T-box gene, Tbx1. Nat Genet. 2001;27:286–291. doi: 10.1038/85845

25. Merscher S, Funke B, Epstein JA, Heyer J, Puech A, Lu MM, Xavier RJ, Demay MB, Russell RG, Factor S, et al. TBX1 is responsible for cardiovascular defects in velo-cardio-facial/DiGeorge syndrome. In: Cell. 2001:619–629.

26. Lindsay EA, Vitelli F, Su H, Morishima M, Huynh T, Pramparo T, Jurecic V, Ogunrinu G, Sutherland HF, Scambler PJ, et al. Tbx1 haploinsufficieny in the DiGeorge syndrome region causes aortic arch defects in mice. Nature. 2001;410:97–101. doi: 10.1038/35065105

27. Anderson RH, Bamforth SD. Morphogenesis of the Mammalian Aortic Arch Arteries. Front Cell Dev Biol. 2022;10:892900. doi: 10.3389/fcell.2022.892900

28. Ramirez A, Astrof S. Visualization and Analysis of Pharyngeal Arch Arteries using Whole-mount Immunohistochemistry and 3D Reconstruction. J Vis Exp. 2020. doi: 10.3791/60797

29. Hutson MR, Kirby ML. Model systems for the study of heart development and disease. Cardiac neural crest and conotruncal malformations. Semin Cell Dev Biol. 2007;18:101–110. doi: 10.1016/j.semcdb.2006.12.004

30. Yagi H, Furutani Y, Hamada H, Sasaki T, Asakawa S, Minoshima S, Ichida F, Joo K, Kimura M, Imamura S-i, et al. Role of TBX1 in human del22q11.2 syndrome. In: The Lancet. 2003:1366–1373.

31. Papangeli I, Scambler PJ. Tbx1 Genetically Interacts With the Transforming Growth Factor-/Bone Morphogenetic Protein Inhibitor Smad7 During Great Vessel Remodeling. In: Circulation Research. 2013:90–102.

32. Zhang Z, Cerrato F, Xu H, Vitelli F, Morishima M, Vincentz J, Furuta Y, Ma L, Martin JF, Baldini A, et al. Tbx1 expression in pharyngeal epithelia is necessary for pharyngeal arch artery development. In: Development (Cambridge, England). 2005:5307–5315.

33. Phillips HM, Stothard CA, Shaikh Qureshi WM, Kousa AI, Briones-Leon JA, Khasawneh RR, O’Loughlin C, Sanders R, Mazzotta S, Dodds R, et al. Pax9 is required for cardiovascular development and interacts with Tbx1 in the pharyngeal endoderm to control 4th pharyngeal arch artery morphogenesis. In: Development. 2019:dev177618.

34. Ryckebusch L, Bertrand N, Mesbah K, Bajolle F, Niederreither K, Kelly RG, Zaffran S. Decreased levels of embryonic retinoic acid synthesis accelerate recovery from arterial growth delay in a mouse model of DiGeorge syndrome. Circ Res. 2010;106:686–694. doi: 10.1161/CIRCRESAHA.109.205732

35. Chen HI, Sharma B, Akerberg BN, Numi HJ, Kivela R, Saharinen P, Aghajanian H, McKay AS, Bogard PE, Chang AH, et al. The sinus venosus contributes to coronary vasculature through VEGFC-stimulated angiogenesis. In: Development. 2014:4500–4512.

36. Apte RS, Chen DS, Ferrara N. VEGF in Signaling and Disease: Beyond Discovery and Development. Cell. 2019;176:1248–1264. doi: 10.1016/j.cell.2019.01.021

37. Freyer L, Hsu CW, Nowotschin S, Pauli A, Ishida J, Kuba K, Fukamizu A, Schier AF, Hoodless PA, Dickinson ME, et al. Loss of Apela Peptide in Mice Causes Low Penetrance Embryonic Lethality and Defects in Early Mesodermal Derivatives. Cell Rep. 2017;20:2116–2130. doi: 10.1016/j.celrep.2017.08.014

38. Baral K, D’Amato G, Kuschel B, Bogan F, Jones BW, Large CL, Whatley JD, Red-Horse K, Sharma B. APJ+ cells in the SHF contribute to the cells of aorta and pulmonary trunk through APJ signaling. Dev Biol. 2023;498:77–86. doi: 10.1016/j.ydbio.2023.04.003

39. Shalaby F, Rossant J, Yamaguchi TP, Gertsenstein M, Wu XF, Breitman ML, Schuh AC. Failure of blood-island formation and vasculogenesis in Flk-1-deficient mice. Nature. 1995;376:62–66. doi: 10.1038/376062a0

40. Shalaby F, Ho J, Stanford WL, Fischer KD, Schuh AC, Schwartz L, Bernstein A, Rossant J. A requirement for Flk1 in primitive and definitive hematopoiesis and vasculogenesis. Cell. 1997;89:981–990. doi: 10.1016/s0092-8674(00)80283-4

41. Abrams MJ, Basinger T, Yuan W, Guo CL, Goentoro L. Self-repairing symmetry in jellyfish through mechanically driven reorganization. Proc Natl Acad Sci U S A. 2015;112:E3365–3373. doi: 10.1073/pnas.1502497112

42. Jelier R, Kruger A, Swoger J, Zimmermann T, Lehner B. Compensatory Cell Movements Confer Robustness to Mechanical Deformation during Embryonic Development. Cell Syst. 2016;3:160–171. doi: 10.1016/j.cels.2016.07.005

43. Sharma B, Ho L, Ford GH, Chen HI, Goldstone AB, Woo YJ, Quertermous T, Reversade B, Red-Horse K. Alternative Progenitor Cells Compensate to Rebuild the Coronary Vasculature in Elabela- and Apj-Deficient Hearts. Dev Cell. 2017;42:655–666 e653. doi: 10.1016/j.devcel.2017.08.008

44. Chen D, Wang X, Liang D, Gordon J, Mittal A, Manley N, Degenhardt K, Astrof S. Fibronectin signals through integrin alpha5beta1 to regulate cardiovascular development in a cell type-specific manner. Dev Biol. 2015;407:195–210. doi: 10.1016/j.ydbio.2015.09.016

45. Liang D, Wang X, Mittal A, Dhiman S, Hou SY, Degenhardt K, Astrof S. Mesodermal expression of integrin alpha5beta1 regulates neural crest development and cardiovascular morphogenesis. Dev Biol. 2014;395:232–244. doi: 10.1016/j.ydbio.2014.09.014

46. van der Flier A, Badu-Nkansah K, Whittaker CA, Crowley D, Bronson RT, Lacy-Hulbert A, Hynes RO. Endothelial alpha5 and alphav integrins cooperate in remodeling of the vasculature during development. Development. 2010;137:2439–2449. doi: 10.1242/dev.049551

47. Milgrom-Hoffman M, Harrelson Z, Ferrara N, Zelzer E, Evans SM, Tzahor E. The heart endocardium is derived from vascular endothelial progenitors. Development. 2011;138:4777–4787. doi: 10.1242/dev.061192

48. Milgrom-Hoffman M, Michailovici I, Ferrara N, Zelzer E, Tzahor E. Endothelial cells regulate neural crest and second heart field morphogenesis. Biol Open. 2014;3:679–688. doi: 10.1242/bio.20148078

49. Matsumoto K, Yoshitomi H, Rossant J, Zaret KS. Liver organogenesis promoted by endothelial cells prior to vascular function. Science. 2001;294:559–563. doi: 10.1126/science.1063889

50. Lammert E, Cleaver O, Melton D. Induction of pancreatic differentiation by signals from blood vessels. Science. 2001;294:564–567. doi: 10.1126/science.1064344

51. Rafii S, Butler JM, Ding BS. Angiocrine functions of organ-specific endothelial cells. Nature. 2016;529:316–325. doi: 10.1038/nature17040

52. Red-Horse K, Ueno H, Weissman IL, Krasnow MA. Coronary arteries form by developmental reprogramming of venous cells. Nature. 2010;464:549–553. doi: 10.1038/nature08873

53. Su T, Stanley G, Sinha R, D’Amato G, Das S, Rhee S, Chang AH, Poduri A, Raftrey B, Dinh TT, et al. Single-cell analysis of early progenitor cells that build coronary arteries. Nature. 2018;559:356–362. doi: 10.1038/s41586-018-0288-7

54. Hou S, Li Z, Dong J, Gao Y, Chang Z, Ding X, Li S, Li Y, Zeng Y, Xin Q, et al. Heterogeneity in endothelial cells and widespread venous arterialization during early vascular development in mammals. Cell Res. 2022;32:333–348. doi: 10.1038/s41422-022-00615-z

55. Xu C, Hasan SS, Schmidt I, Rocha SF, Pitulescu ME, Bussmann J, Meyen D, Raz E, Adams RH, Siekmann AF. Arteries are formed by vein-derived endothelial tip cells. In: Nature Communications. 2014:5758.

56. Lee HW, Xu Y, He L, Choi W, Gonzalez D, Jin SW, Simons M. Role of Venous Endothelial Cells in Developmental and Pathologic Angiogenesis. Circulation. 2021;144:1308–1322. doi: 10.1161/CIRCULATIONAHA.121.054071

57. Park H, Furtado J, Poulet M, Chung M, Yun S, Lee S, Sessa WC, Franco CA, Schwartz MA, Eichmann A. Defective Flow-Migration Coupling Causes Arteriovenous Malformations in Hereditary Hemorrhagic Telangiectasia. Circulation. 2021;144:805–822. doi: 10.1161/CIRCULATIONAHA.120.053047

58. Bussmann J, Wolfe SA, Siekmann AF. Arterial-venous network formation during brain vascularization involves hemodynamic regulation of chemokine signaling. Development. 2011;138:1717–1726. doi: 10.1242/dev.059881

59. Hasan SS, Tsaryk R, Lange M, Wisniewski L, Moore JC, Lawson ND, Wojciechowska K, Schnittler H, Siekmann AF. Endothelial Notch signalling limits angiogenesis via control of artery formation. Nat Cell Biol. 2017;19:928–940. doi: 10.1038/ncb3574

60. Fujita M, Cha YR, Pham VN, Sakurai A, Roman BL, Gutkind JS, Weinstein BM. Assembly and patterning of the vascular network of the vertebrate hindbrain. Development. 2011;138:1705–1715. doi: 10.1242/dev.058776

61. Paulissen SM, Castranova DM, Krispin SM, Burns MC, Menendez J, Torres-Vazquez J, Weinstein BM. Anatomy and development of the pectoral fin vascular network in the zebrafish. Development. 2022;149. doi: 10.1242/dev.199676

62. Leonard EV, Hasan SS, Siekmann AF. Temporally and regionally distinct morphogenetic processes govern zebrafish caudal fin blood vessel network expansion. Development. 2023;150. doi: 10.1242/dev.201030

63. Cai CL, Liang X, Shi Y, Chu PH, Pfaff SL, Chen J, Evans S. Isl1 identifies a cardiac progenitor population that proliferates prior to differentiation and contributes a majority of cells to the heart. Dev Cell. 2003;5:877–889. doi: 10.1016/s1534-5807(03)00363-0

64. Evans SM, Yelon D, Conlon FL, Kirby ML. Myocardial lineage development. Circ Res. 2010;107:1428-1444. doi: 10.1161/CIRCRESAHA.110.227405

65. Sun Y, Liang X, Najafi N, Cass M, Lin L, Cai CL, Chen J, Evans SM. Islet 1 is expressed in distinct cardiovascular lineages, including pacemaker and coronary vascular cells. Dev Biol. 2007;304:286–296. doi: 10.1016/j.ydbio.2006.12.048

66. Devine WP, Wythe JD, George M, Koshiba-Takeuchi K, Bruneau BG. Early patterning and specification of cardiac progenitors in gastrulating mesoderm. Elife. 2014;3. doi: 10.7554/eLife.03848

67. Verzi MP, McCulley DJ, De Val S, Dodou E, Black BL. The right ventricle, outflow tract, and ventricular septum comprise a restricted expression domain within the secondary/anterior heart field. Dev Biol. 2005;287:134–145. doi: 10.1016/j.ydbio.2005.08.041

68. Zhang Z, Huynh T, Baldini A. Mesodermal expression of Tbx1 is necessary and sufficient for pharyngeal arch and cardiac outflow tract development. Development. 2006;133:3587–3595. doi: 10.1242/dev.02539

69. Verzi MP, McCulley DJ, De Val S, Dodou E, Black BL. The right ventricle, outflow tract, and ventricular septum comprise a restricted expression domain within the secondary/anterior heart field. In: Developmental Biology. 2005:134–145.

70. Ema M, Takahashi S, Rossant J. Deletion of the selection cassette, but not cis-acting elements, in targeted Flk1-lacZ allele reveals Flk1 expression in multipotent mesodermal progenitors. In: Blood. 2006:111–117.

71. Hooper AT, Butler JM, Nolan DJ, Kranz A, Iida K, Kobayashi M, Kopp HG, Shido K, Petit I, Yanger K, et al. Engraftment and Reconstitution of Hematopoiesis Is Dependent on VEGFR2-Mediated Regeneration of Sinusoidal Endothelial Cells. In: Cell Stem Cell. 2009:263–274.

72. Madisen L, Zwingman TA, Sunkin SM, Oh SW, Zariwala HA, Gu H, Ng LL, Palmiter RD, Hawrylycz MJ, Jones AR, et al. A robust and high-throughput Cre reporting and characterization system for the whole mouse brain. Nat Neurosci. 2010;13:133–140. doi: 10.1038/nn.2467

73. Muzumdar MD, Tasic B, Miyamichi K, Li L, Luo L. A global double-fluorescent Cre reporter mouse. Genesis. 2007;45:593–605. doi: 10.1002/dvg.20335

74. Yang L, Cai CL, Lin L, Qyang Y, Chung C, Monteiro RM, Mummery CL, Fishman GI, Cogen A, Evans S. Isl1Cre reveals a common Bmp pathway in heart and limb development. Development. 2006;133:1575–1585. doi: 10.1242/dev.02322

75. Arnold SM, Fessler LI, Fessler JH, Kaufman RJ. Two homologues encoding human UDP-glucose:glycoprotein glucosyltransferase differ in mRNA expression and enzymatic activity. Biochemistry. 2000;39:2149–2163. doi: 10.1021/bi9916473

76. Sousa VH, Miyoshi G, Hjerling-Leffler J, Karayannis T, Fishell G. Characterization of Nkx6-2-derived neocortical interneuron lineages. Cereb Cortex. 2009;19 Suppl 1:i1–10. doi: 10.1093/cercor/bhp038

77. Madisen L, Garner AR, Shimaoka D, Chuong AS, Klapoetke NC, Li L, van der Bourg A, Niino Y, Egolf L, Monetti C, et al. Transgenic Mice for Intersectional Targeting of Neural Sensors and Effectors with High Specificity and Performance. In: Neuron. 2015:942–958.

78. Ehlers ML, Celona B, Black BL. NFATc1 controls skeletal muscle fiber type and is a negative regulator of MyoD activity. Cell Rep. 2014;8:1639–1648. doi: 10.1016/j.celrep.2014.08.035

79. Sinha T, Lammerts van Bueren K, Dickel DE, Zlatanova I, Thomas R, Lizama CO, Xu SM, Zovein AC, Ikegami K, Moskowitz IP, et al. Differential Etv2 threshold requirement for endothelial and erythropoietic development. Cell Rep. 2022;39:110881. doi: 10.1016/j.celrep.2022.110881

80. Astrof S, Kirby A, Lindblad-Toh K, Daly M, Hynes RO. Heart development in fibronectin-null mice is governed by a genetic modifier on chromosome four. Mech Dev. 2007;124:551–558. doi: 10.1016/j.mod.2007.05.004

81. Devine WP, Wythe JD, George M, Koshiba-Takeuchi K, Bruneau BG. Early patterning and specification of cardiac progenitors in gastrulating mesoderm. In: eLife. 2014:1–23.

82. Liu Z, Chen O, Wall JBJ, Zheng M, Zhou Y, Wang L, Vaseghi HR, Qian L, Liu J. Systematic comparison of 2A peptides for cloning multi-genes in a polycistronic vector. Sci Rep. 2017;7:2193. doi: 10.1038/s41598-017-02460-2

83. Nomaru H, Liu Y, De Bono C, Righelli D, Cirino A, Wang W, Song H, Racedo SE, Dantas AG, Zhang L, et al. Single cell multi-omic analysis identifies a Tbx1-dependent multilineage primed population in murine cardiopharyngeal mesoderm. Nat Commun. 2021;12:6645. doi: 10.1038/s41467-021-26966-6

84. Schindelin J, Arganda-Carreras I, Frise E, Kaynig V, Longair M, Pietzsch T, Preibisch S, Rueden C, Saalfeld S, Schmid B, et al. Fiji: an open-source platform for biological-image analysis. Nat Methods. 2012;9:676–682. doi: 10.1038/nmeth.2019

85. Lord SJ, Velle KB, Mullins RD, Fritz-Laylin LK. SuperPlots: Communicating reproducibility and variability in cell biology. J Cell Biol. 2020;219. doi: 10.1083/jcb.202001064

86. Morse A. Formic Acid-Sodium Citrate Decalcification and Butyl Alcohol Dehydration of Teeth and Bones for Sectioning in Paraffin. Journal of Dental Research. 1945:143–153. doi: 10.1177

87. Degenhardt K, Wright AC, Horng D, Padmanabhan A, Epstein JA. Rapid 3D Phenotyping of Cardiovascular Development in Mouse Embryos by Micro-CT With Iodine Staining. In: Circulation: Cardiovascular Imaging. 2010:314–322.

88. Ema M, Takahashi S, Rossant J. Deletion of the selection cassette, but not cis-acting elements, in targeted Flk1-lacZ allele reveals Flk1 expression in multipotent mesodermal progenitors. Blood. 2006;107:111–117. doi: 10.1182/blood-2005-05-1970

89. Arnold JS, Werling U, Braunstein EM, Liao J, Nowotschin S, Edelmann W, Hebert JM, Morrow BE. Inactivation of Tbx1 in the pharyngeal endoderm results in 22q11DS malformations. Development. 2006;133:977–987. doi: 10.1242/dev.02264

