## Supplemental Figures for "Identification of novel buffering mechanisms in aortic arch artery development and congenital heart disease"

Supplemental Figure 1

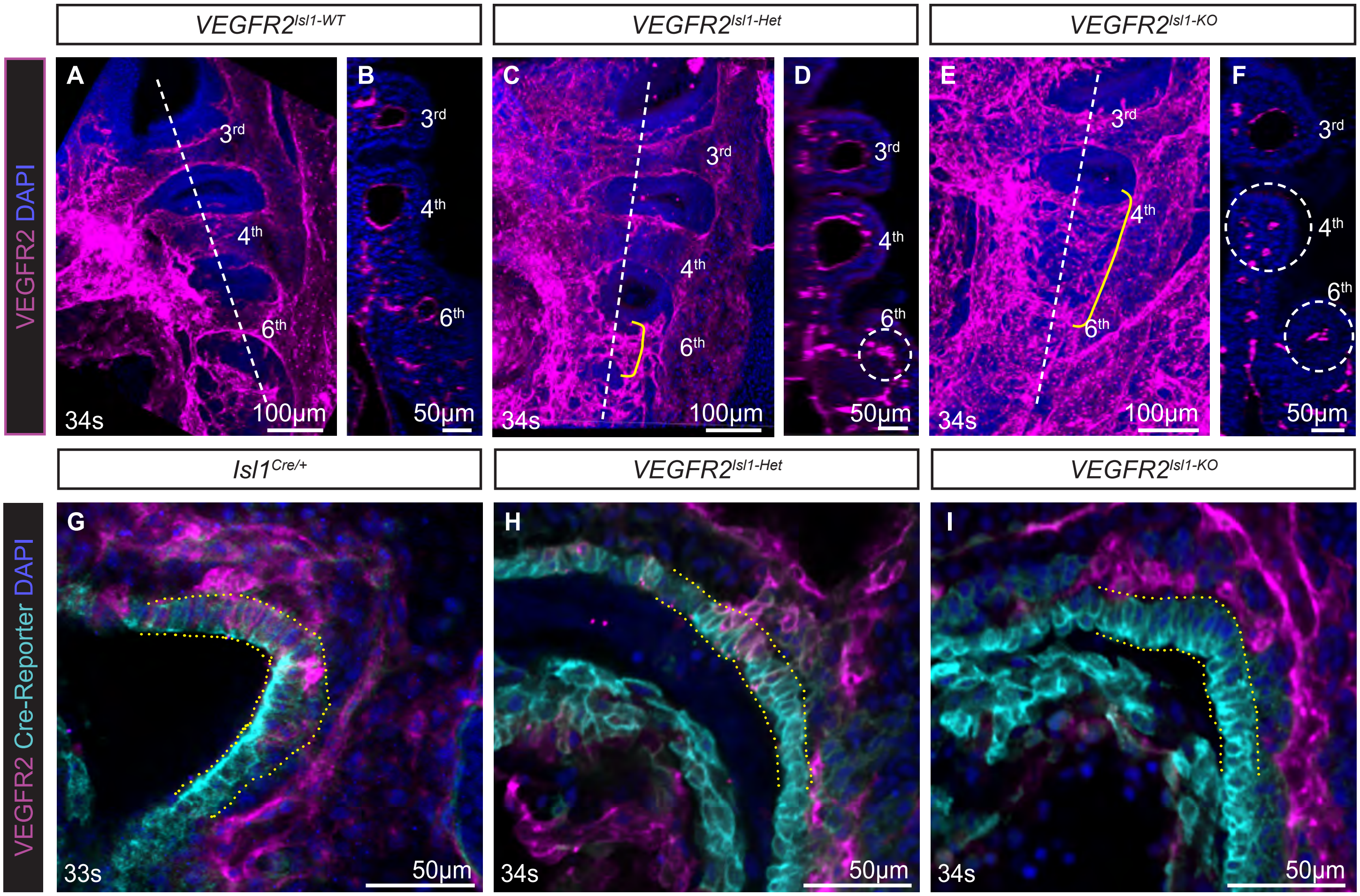


**Figure S1. VEGFR2 is required in the *Isl1*-lineage for PAA development.**
**A-F,** Whole E10.5 embryos were stained by immunofluorescence and imaged by confocal microscopy to detect VEGFR2. Maximum intensity projections through the pharyngeal region are shown in **A, C** and **E**. Pharyngeal arches are numbered. Vertical dashed lines mark the planes of sections shown in **B, D,** and **F**. Yellow bracket in **C** marks malformed 6th PAA. Dashed circles in **D** and **F** mark 4th and 6th pharyngeal arches. Note the presence of EC plexus rather than a large central 4th and 6th PAAs. **G-I.** Sagittal optical sections through E10.5 embryos stained for VEGFR2 and Isl1-lineage marker. Note the downregulation of VEGFR2 in the SHF-derived cells of the dorsal pericardial wall in *VEGFR2*$^{Isl1\text{-}KO}$ embryos (outlined), **I**.

Supplemental Figure 2

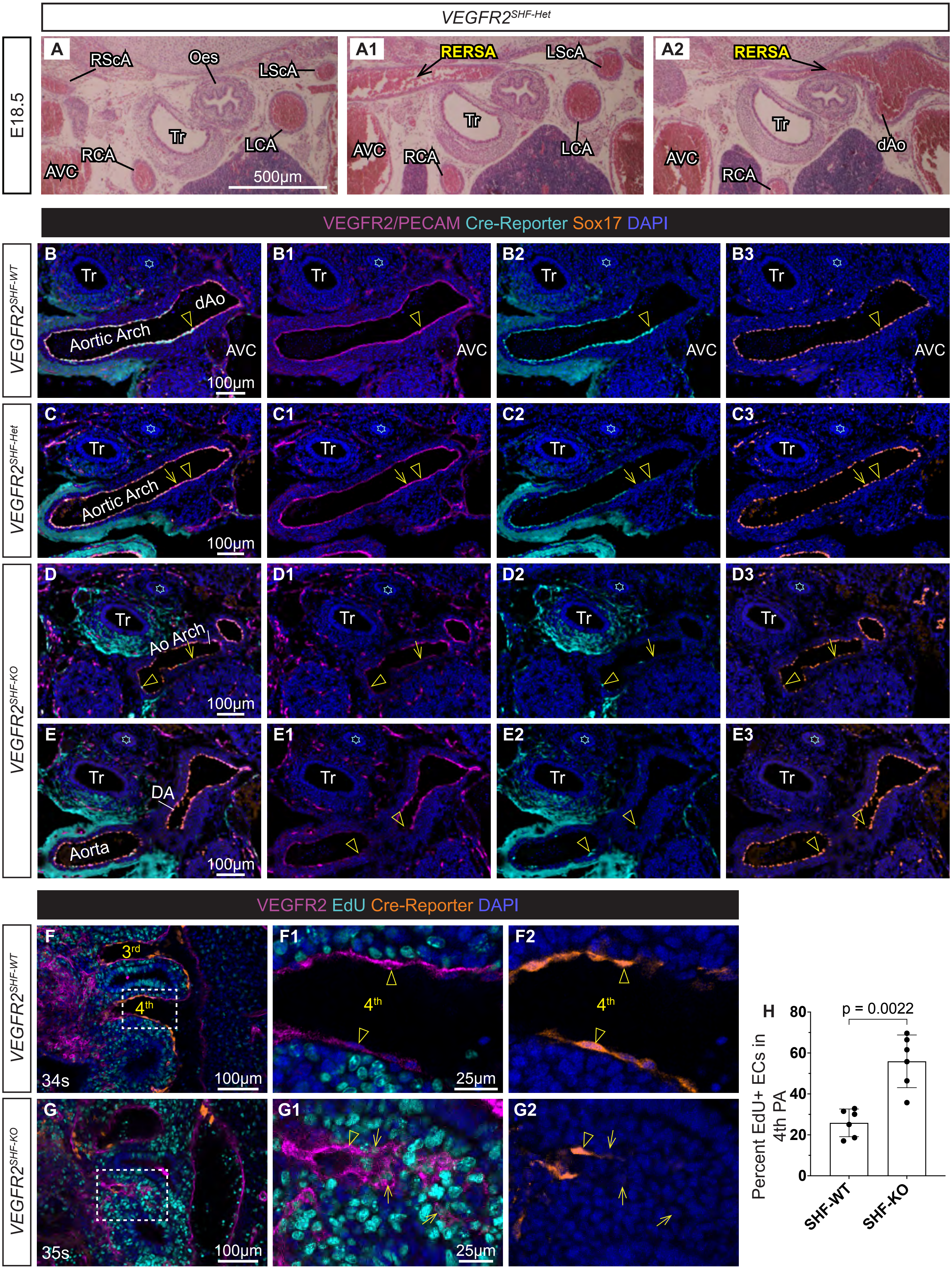

**Figure S2.**

**A, One case of arch artery defects in VEGFR2$^{SHF\text{-}Het}$ embryo: embryo 21, Table S6.** Transverse sections of E18.5 *VEGFR2*$^{SHF\text{-}Het}$ embryo showing RERSA (arrow) stained with hematoxylin and eosin.

**B-E3. ECs from the alternative source incorporate into the mature aortic arch and its arteries in *VEGFR2*$^{SHF\text{-}Het}$ and *VEGFR2*$^{SHF\text{-}KO}$ embryos.** Confocal 0.45 μm images of transverse frozen tissue sections from E14.5 embryos were stained to detect Pecam-1 in *VEGFR2*$^{SHF\text{-}WT}$ or by detecting GFP expressed from the Flk1-GFP allele in lieu of VEGFR2 (see Mice, in Methods; magenta), SHF lineage (Cre-reporter, cyan), Sox17 (orange) and nuclei (DAPI, blue). Arrowheads point to SHF-lineage cells. **B-B3,** *VEGFR2*$^{SHF\text{-}WT}$ embryo. Most of the aortic arch ECs are SHF-derived. Cre-reporter expression is absent in the descending aorta (dAo). Note the absence of Sox17$^{+}$ ECs in the anterior vena cava (AVC). Tr-trachea. Blue stars mark esophagus in each section. **C-C3,** *VEGFR2*$^{SHF\text{-}Het}$ embryo. Arrows point to non-SHF-lineage cells. **D-E,** Most ECs in the aortic arch, ascending aorta and ductus in *VEGFR2*$^{SHF\text{-}KO}$ embryo are derived from a non-SHF source. **D-D3,** Sections through the aortic arch. **E-E3,** Sections through the ascending aorta and ductus arteriosus (DA). Arrowheads point to the rare SHF-derived cells. Arrows point to non-SHF-lineage cells. Note that the majority of ECs in the aortic arch, ascending aorta and ductus in *VEGFR2*$^{SHF\text{-}KO}$ embryos express an arterial marker Sox17.

**F-G, Increased proliferation of pharyngeal arch ECs in *VEGFR2*$^{SHF\text{-}KO}$ embryos.** Pregnant females were injected with 125 μg of EdU 30 min prior to dissection at E10.5. Whole embryos were stained to detect VEGFR2 (magenta), Cre-reporter (orange), and EdU (cyan). 0.45 μm sagittal optical sections are shown. Regions of the 4th arches in **F** and **G** are expanded to the right. Note that the majority of PAA ECs in the 4th PAA of *VEGFR2*$^{SHF\text{-}WT}$ control are EdU-negative (arrowheads), **F1-F2**. In *VEGFR2*$^{SHF\text{-}KO}$ mutant, arrowhead points to EdU-negative SHF-lineage cell and arrows point to multiple EdU+ com-

Supplemental Figure 3

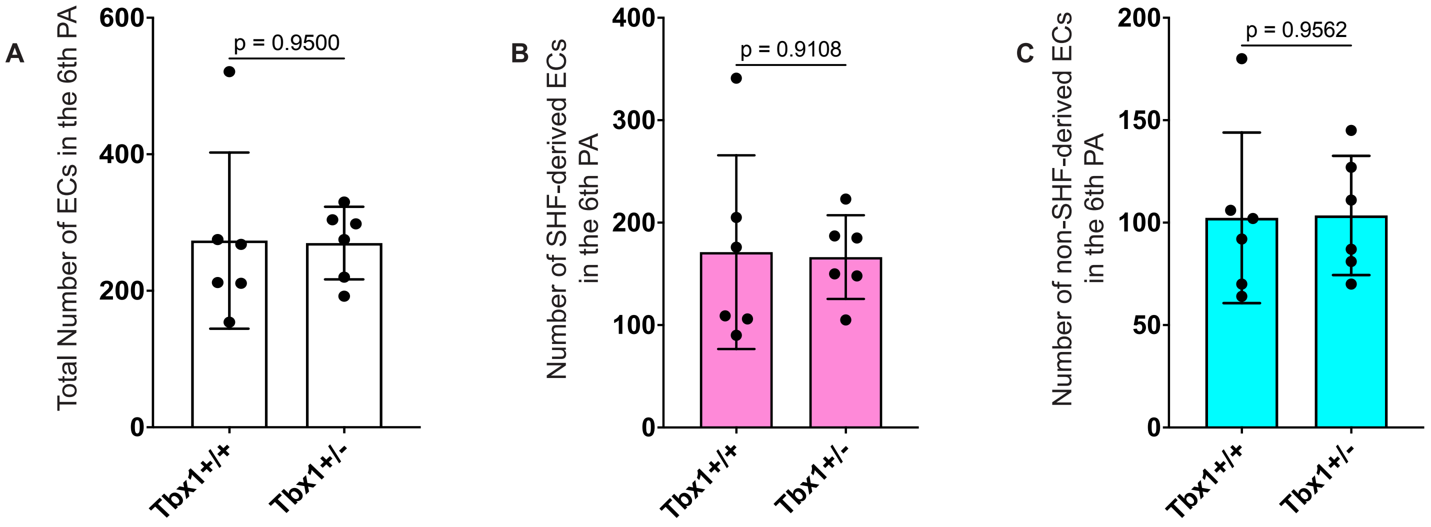


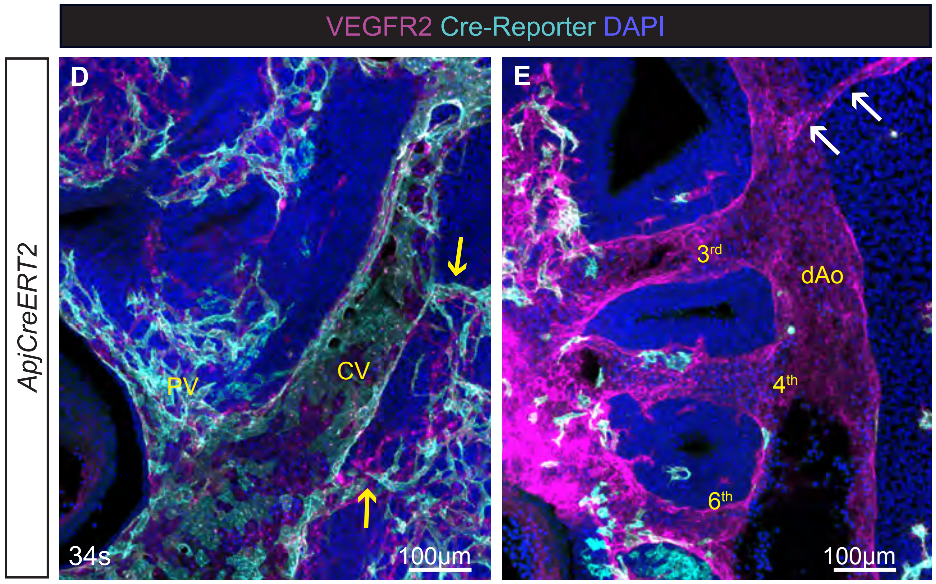


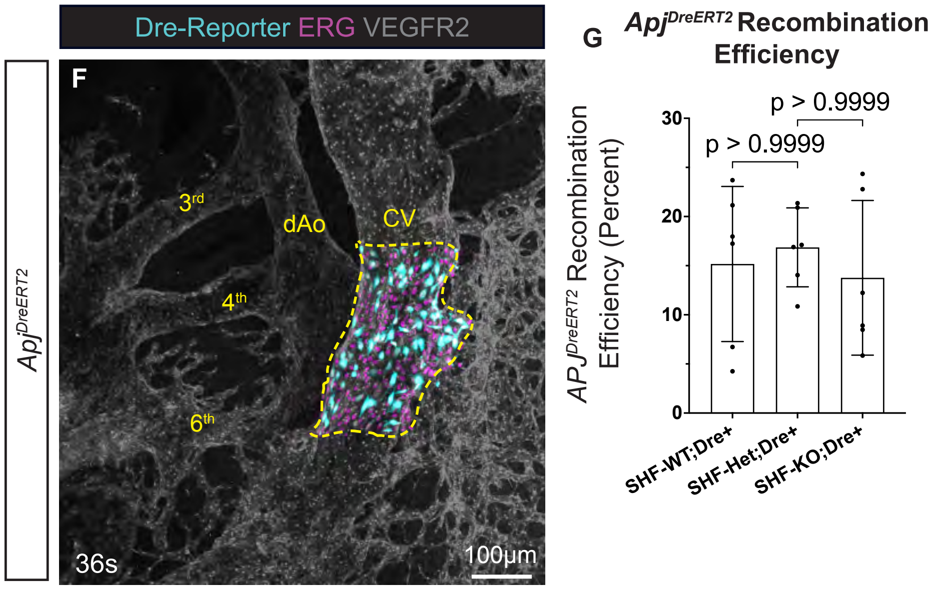


**Figure S3.**
**A-C, Heterozygocity in *Tbx1* does not affect EC composition of the 6th pharyngeal arch.** Quantification of endothelial populations in the 6th pharyngeal arches of *Tbx1*$^{+/-}$ embryos and their control litter mates at E10.5. Each dot represents one arch. Mean and standard deviations are shown, p values were calculated using 2-tailed, unpaired, non-parametric Mann-Whitney test.

**D-E**, **Vein labeling using Apj-CreERT2 line.** Tamoxifen injection at E9.5. Whole E10.5 embryos were stained by immunofluorescence (IF) and imaged by confocal microscopy to detect VEGFR2 (magenta) and Cre-reporter (cyan). **D**, Maximum intensity projection through 50µm-thick sagittal optical sections show *Apj-CreERT2* lineage labeling in the cardinal vein (CV) and intersomitic veins (yellow arrows) but not in the dAo or intersomitic arteries (white arrows), **E**. Few Cre-reporter cells can be seen in the PAA endothelium. PAAs are numbered. dAo - dorsal aorta.

Supplemental Figure 4

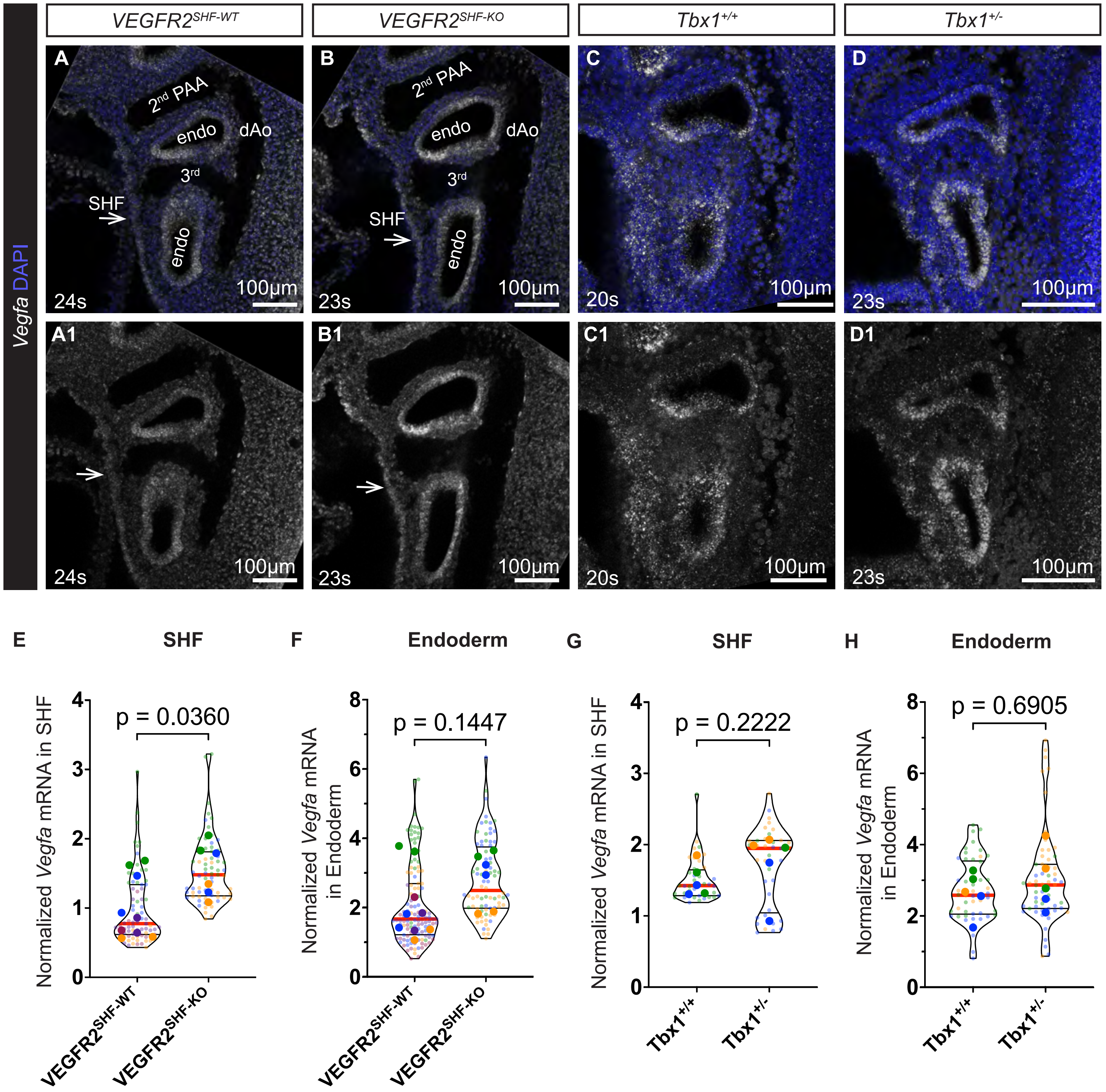


**Figure S4. Loss of SHF-derived vascular progenitors increases *VEGFA* mRNA expression in the in *VEGFR2$^{SHF-KO}$* embryos but not in *Tbx1$^{+/-}$* embryos.**

Single-molecule RNA fluorescent *in situ* hybridization of E9.5 embryos to detect the expression of *VEGF-A* (white). **A,A1,** *VEGFR2$^{SHF-WT}$* embryos. **B, B1,** *VEGFR2$^{SHF-KO}$* embryos. **C,C1,** *Tbx1$^{+/+}$* embryos. **D, D1,** *Tbx1$^{+/-}$* embryos. **E-H,** Normalized VEGFA mRNA intensity was determined by dividing mean fluorescence intensity in the SHF or pharyngeal endoderm by the mean fluorescence intensity in the neural tube of the embryo in the same optical section. Light-colored dots mark individual measurements in serial optical sections. Dark-colored dots mark the average of measurements per pharyngeal arch. Red lines mark means, and black lines mark quartiles. *VEGFR2$^{SHF-WT}$* n = 6 arches, *VEGFR2$^{SHF-KO}$* n = 6 arches, *Tbx1$^{+/+}$* n=5 arches, *Tbx1$^{+/-}$* n=5 arches. Mean values (large dots) from each experiment were used in 2-tailed, unpaired, non-parametric Mann-Whitney 2-tailed t tests to calculate p values.

Supplemental Figure 5

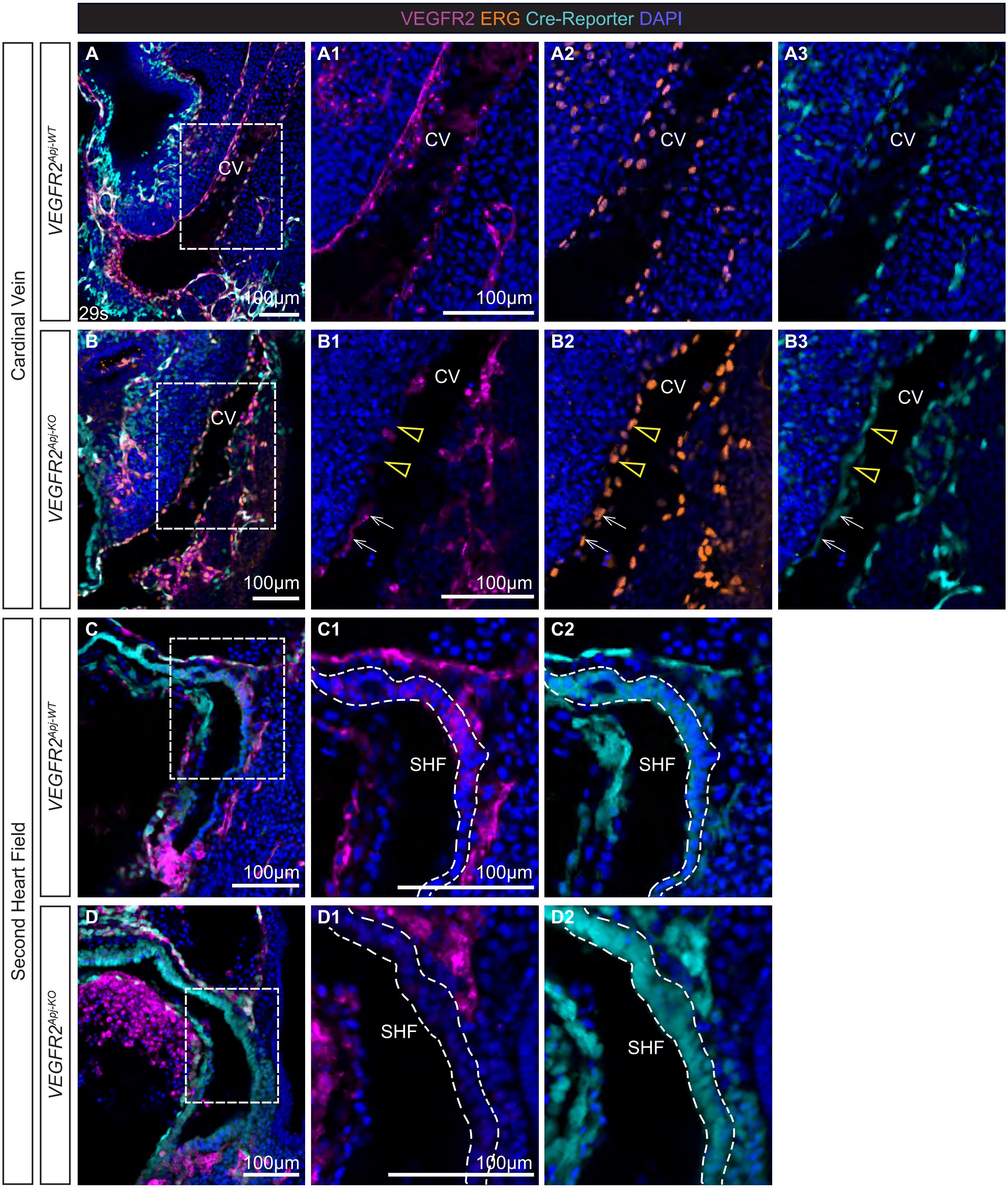

**Figure S5. Tamoxifen injection at E8.5 leads to efficient labeling of SHF- and vein-derived cells, and the ablation of VEGFR2 in these lineages in Apj-CreERT2 strain.** Following injection of tamoxifen at E8.5, VEGFR2$^{Apj\text{-}WT}$ and VEGFR2$^{Apj\text{-}KO}$ embryos were isolated at E10.0 and stained whole using immunofluorescence (IF) to detect VEGFR2 (magenta), ERG (orange), Cre-reporter (cyan), and nuclei (DAPI, blue). Confocal microscopy was used to image embryos, 5 μm-thick sagittal optical sections are shown. Boxed regions in **A-D** are expanded in panels to the right. **A-B,** Regions containing the cardinal veins (CV) are shown. **A-A3**, VEGFR2$^{Apj\text{-}WT}$. **B-B3,** VEGFR2$^{Apj\text{-}KO}$. Note VEGFR2-negative ECs (arrowheads) and VEGFR2+ escapees (white arrows) in the CV of VEGFR2$^{Apj\text{-}KO}$ mutant, **B1-B3**. **C-D,** Regions showing SHF-derived dorsal pericardial wall (outlined by the dashed lines in **C1-C2** and **D1-D2** are shown. Note the absence of VEGFR2 expression in the SHF of VEGFR2$^{Apj\text{-}KO}$ mutant, **D1-D2**.
