## Supplemental Tables for "Identification of novel buffering mechanisms in aortic arch artery development and congenital heart disease"

### Table S1

| Genotype | n = arches | 3rd PAA | | 4th PAA | | 6th PAA | |
|---|---|---|---|---|---|---|---|
| | | Hypoplastic | Aplastic | Hypoplastic | Aplastic | Hypoplastic | Aplastic |
| **VEGFR2$^{Isl1\text{-}WT}$**<br>*VEGFR2$^{flox/-}$; Isl1Cre$^{Cre/+}$* | 5 | 0 | 0 | 0 | 0 | 0 | 0 |
| **VEGFR2$^{Isl1\text{-}HET}$**<br>*VEGFR2$^{flox/-}$; Isl1Cre$^{Cre/+}$* | 7 | 0 | 0 | 0 | 0 | 0 | 1<br>(14%) |
| **VEGFR2$^{Isl1\text{-}KO}$**<br>*VEGFR2$^{flox/-}$; Isl1Cre$^{Cre/+}$* | 5 | 0 | 0 | 1<br>(20%) | 4<br>(80%) | 1<br>(20%) | 4<br>(80%) |
| **VEGFR2$^{SHF\text{-}WT}$**<br>*VEGFR2$^{+/+}$;Mef2c-AHF-Cre* | 10 | 0 | 0 | 0 | 0 | 0 | 0 |
| **VEGFR2$^{SHF\text{-}Het}$**<br>*VEGFR2$^{flox/+}$;Mef2c-AHF-Cre* | 11 | 0 | 0 | 1<br>(9.1%) | 0 | 3<br>(27.3%) | 0 |
| **VEGFR2$^{SHF\text{-}KO}$**<br>*VEGFR2$^{flox/-}$;Mef2c-AHF-Cre* | 12 | 0 | 0 | 3<br>(25%) | 9<br>(75%) | 0 | 12<br>(100%) |
| **Tbx1$^{+/+}$**<br>*Tbx1$^{+/+}$;Mef2c-AHF-Cre* | 6 | 0 | 0 | 0 | 0 | 0 | 0 |
| **Tbx1$^{+/-}$**<br>*Tbx1$^{+/-}$;Mef2c-AHF-Cre* | 6 | 0 | 0 | 3<br>(50%) | 3<br>(50%) | 1<br>(16.7%) | 1<br>(16.7%) |
| **Tbx1$^{SHF\text{-}Het}$**<br>*Tbx1$^{flox/+}$;Mef2c-AHF-Cre* | 6 | 0 | 0 | 0 | 0 | 0 | 0 |
| **Tbx1$^{SHF\text{-}Null}$**<br>*Tbx1$^{flox/flox}$;Mef2c-AHF-Cre* | 8 | 1 | 0 | 1<br>(12.5%) | 7<br>(87.5%) | 5<br>(62.5%) | 1<br>(12.5%) |

**Supplemental Table 1. Penetrance of aberrant PAA formation.** PAAs of E10.5 embryos from 34-36s were analyzed by whole-mount immunofluorescence (WM-IF) staining for VEGFR2. Aberrant PAAs were marked as “hypoplastic” when their diameter was markedly smaller than controls (e.g., **Fig. 2C2**) or as “aplastic,” indicating the absence of a central PAA lumen (e.g., **Fig. 2E2**). “0” indicates PAAs comparable to controls. n = the number of individual pharyngeal arches analyzed. Percent of penetrance is reported in parentheses.

### Table S2

| Genotype | Pharyngeal Arch | Number of SHF-derived cells (Cre-reporter+) | Percent of VEGFR2+ SHF cells (VEGFR2+, Cre-reporter+) | Percent of VEGFR2- SHF cells (VEGFR2-, Cre-reporter+) |
|---|---|---|---|---|
| **VEGFR2$^{SHF\text{-}WT}$** *VEGFR2$^{+/+}$;Mef2c-AHF-Cre* (n = 8 arches) | 3rd | 279 ± 89 | 65.3% | 34.7% |
| | 4th | 501 ± 133 | 65.5% | 34.5% |
| | 6th | 357 ± 86 | 80.0% | 20.0% |
| **VEGFR2$^{SHF\text{-}KO}$** *VEGFR2$^{flox/-}$;Mef2c-AHF-Cre* (n = 6 arches) | 3rd | 267 ± 46 | 16.0% | 84.0% |
| | 4th | 373 ± 164 | 11.8% | 88.2% |
| | 6th | 113 ± 68 | 11.9% | 88.1% |

**Supplemental Table 2. The average number of SHF-derived cells in the pharyngeal arches.** Pharyngeal arches of E10.5 *VEGFR2$^{SHF\text{-}WT}$* and *VEGFR2$^{SHF\text{-}KO}$* embryos ranging from 34-36s were analyzed via WM-IF staining. The total number of SHF cells was quantified in each pharyngeal arch. Percent of VEGFR2-positive and VEGFR2-negative SHF cells was determined. Averages ± SD are reported.

### Table S3

| Genotype | Pharyngeal Arch | Number of ECs (VEGFR2+, ERG+) | Percent SHF-derived ECs (VEGFR2+, ERG+, Cre-reporter+) | Percent Non-SHF-derived ECs (VEGFR+, ERG+, Cre-reporter-) |
|---|---|---|---|---|
| **VEGFR2$^{SHF-WT}$** *VEGFR2$^{+/+}$;Mef2c-AHF-Cre* (n = 10 arches) | 3rd | 508 ± 120 | 36.1 ± 7.4% | 63.9 ± 7.4% |
| | 4th | 476 ± 81 | 71.6 ± 8.8% | 28.4 ± 8.8% |
| | 6th | 455 ± 143 | 69.2 ± 11.9% | 30.8 ± 11.9% |
| **VEGFR2$^{SHF-Het}$** *VEGFR2$^{flox/+}$;Mef2c-AHF-Cre* (n = 11 arches) | 3rd | 520 ± 110 | 21.9 ± 6.7% | 78.1 ± 6.7% |
| | 4th | 567 ± 72 | 41.2 ± 5.9% | 58.8 ± 5.9% |
| | 6th | 406 ± 132 | 39.0 ± 9.1% | 61.0 ± 9.1% |
| **VEGFR2$^{SHF-KO}$** *VEGFR2$^{flox/-}$;Mef2c-AHF-Cre* (n = 12 arches) | 3rd | 517 ± 106 | 10.1 ± 5.6% | 89.8 ± 5.6% |
| | 4th | 585 ± 114 | 6.3 ± 3.1% | 93.7 ± 3.1% |
| | 6th | 350 ± 138 | 3.5 ± 1.6% | 96.5 ± 1.6% |

**Supplemental Table 3. The average number of ECs in the pharyngeal arches.** Pharyngeal arches of E10.5 *VEGFR2$^{SHF-WT}$*, *VEGFR2$^{SHF-Het}$*, and *VEGFR2$^{SHF-KO}$* embryos ranging from 34-36s were analyzed via WM-IF for VEGFR2, Cre-reporter, and ERG. The total number of EC cells. percent of SHF-derived ECs and percent of non-SHF-derived ECs were quantified. Averages ± SD are reported. Statistical analyses were reported in **Fig. 2H-I**.

### Table S4

| Genotype | Pharyngeal Arch | Number of SHF-derived ECs (ERG+, Cre-reporter+) | Percent of SHF-derived ECs that are VEGFR2+ (VEGFR2+, ERG+, Cre-reporter+) | Percent VEGFR2- Cre-reporter+ ECs (VEGFR2-, ERG+, Cre-reporter+) |
|---|---|---|---|---|
| **VEGFR2$^{SHF\text{-}KO}$** *VEGFR2$^{flox/-}$;Mef2c-AHF-Cre* (n = 10 arches) | $3^{rd}$ | 57 ± 33 | 84.7 ± 14.4% | 14.3 ± 14.4% |
| | $4^{th}$ | 37 ± 15 | 87.6 ± 15.5% | 12.4 ± 15.5% |
| | $6^{th}$ | 11 ± 5 | 98.3 ± 3.8% | 1.7 ± 3.8% |

**Supplemental Table 4. Cells that have escaped recombination at the VEGFR2 locus in VEGFR2$^{SHF\text{-}KO}$ embryos (escapees).** VEGFR2+ SHF-derived ECs in VEGFR2$^{SHF\text{-}KO}$ embryos are cells in which Cre-mediated recombination took place at the ROSA locus but not at the VEGFR2 locus, resulting in the expression of VEGFR2 in Cre-reporter+ cells. Such cells are a minority (~10%) of total pharyngeall arch ECs (see **Fig. 2H**) in VEGFR2$^{SHF\text{-}KO}$ embryos.

### Table S5

**Cardiovascular defects in *VEGFR2*$^{Isl1\text{-}KO}$ and *VEGFR2*$^{SHF\text{-}KO}$ embryos**

| Embryo | Cre-strain | Age | Analysis Type | Arch Artery Defect |
|---|---|---|---|---|
| 1 | *Isl1*$^{Cre}$ | 14.5 | H&E | IAA-B*, Aberrant RScA |
| 2 | *Isl1*$^{Cre}$ | 14.5 | H&E | Aberrant RScA |
| 3 | *Isl1*$^{Cre}$ | 14.5 | H&E | Vascular Ring |
| 4 | *Isl1*$^{Cre}$ | 14.5 | H&E | Aberrant RScA |
| 5 | *Isl1*$^{Cre}$ | 14.5 | H&E | Aberrant RScA and LScA |
| 6 | *Isl1*$^{Cre}$ | 18.5 | μCT | IAA-B*, Aberrant RScA |
| 7 | *Isl1*$^{Cre}$ | 18.5 | μCT | None |
| 8 | *Isl1*$^{Cre}$ | 18.5 | μCT | Vascular Ring, Aberrant LScA |
| 9 | *Isl1*$^{Cre}$ | 18.5 | μCT | IAA-B*, Aberrant RScA |
| 10 | *Isl1*$^{Cre}$ | 18.5 | μCT | Aberrant LScA |
| 11 | *Isl1*$^{Cre}$ | 18.5 | μCT | Aberrant LScA |
| 12 | *Isl1*$^{Cre}$ | 18.5 | μCT | Aberrant RScA |
| 13 | *Isl1*$^{Cre}$ | 18.5 | μCT | Aberrant RScA |
| 14 | *Isl1*$^{Cre}$ | 18.5 | μCT | None |
| 15 | *Isl1*$^{Cre}$ | 18.5 | μCT | None |
| 16 | *Mef2c-AHF-Cre* | 14.5 | Vibratome | IAA-B* |
| 17 | *Mef2c-AHF-Cre* | 14.5 | Vibratome | IAA-B*, Aberrant RScA |
| 18 | *Mef2c-AHF-Cre* | 14.5 | Vibratome | Cervical aortic arch, pulmonary atresia |
| 19 | *Mef2c-AHF-Cre* | 18.5 | H&E | Aberrant RScA |
| 20 | *Mef2c-AHF-Cre* | 18.5 | H&E | Aberrant RScA |
| 21 | *Mef2c-AHF-Cre* | 18.5 | H&E | IAA-C* |
| 22 | *Mef2c-AHF-Cre* | 18.5 | H&E | Aberrant RScA |
| 23 | *Mef2c-AHF-Cre* | 18.5 | H&E | None |
| **Summary** | | | | **19/23 (82.6%)** |

**Supplemental Table 5. Cardiovascular defects in *VEGFR2*$^{Isl1\text{-}KO}$ and *VEGFR2*$^{SHF\text{-}KO}$ embryos.** **Abbreviations**: H&E – hematoxylin and eosin; μCT – micro-computed tomography; RScA – right subclavian artery; IAA-B – interrupted aortic arch type B; IAA-C – interrupted aortic arch type C. Arch artery defects lethal postnatally are labeled in red and marked by an asterisk.

### Table S6

**Cardiovascular defects in *VEGFR2*$^{Isl1\text{-}Het}$ and *VEGFR2*$^{SHF\text{-}Het}$ embryos**

| Embryo | Cre-strain | Age | Analysis Type | Arch Artery Defect |
|---|---|---|---|---|
| 1 | $Isl1^{Cre}$ | 14.5 | H&E | None |
| 2 | $Isl1^{Cre}$ | 14.5 | H&E | None |
| 3 | $Isl1^{Cre}$ | 18.5 | µCT | None |
| 4 | $Isl1^{Cre}$ | 18.5 | µCT | None |
| 5 | $Isl1^{Cre}$ | 18.5 | µCT | None |
| 6 | $Isl1^{Cre}$ | 18.5 | µCT | None |
| 7 | $Isl1^{Cre}$ | 18.5 | µCT | None |
| 8 | $Isl1^{Cre}$ | 18.5 | µCT | None |
| 9 | $Isl1^{Cre}$ | 18.5 | µCT | None |
| 10 | $Isl1^{Cre}$ | 18.5 | µCT | None |
| 11 | $Isl1^{Cre}$ | 18.5 | µCT | None |
| 12 | $Isl1^{Cre}$ | 18.5 | µCT | None |
| 13 | $Isl1^{Cre}$ | 18.5 | µCT | None |
| 14 | *Mef2c-AHF-Cre* | 14.5 | Vibratome | None |
| 15 | *Mef2c-AHF-Cre* | 14.5 | Vibratome | None |
| 16 | *Mef2c-AHF-Cre* | 14.5 | Vibratome | None |
| 17 | *Mef2c-AHF-Cre* | 18.5 | H&E | None |
| 18 | *Mef2c-AHF-Cre* | 18.5 | H&E | None |
| 19 | *Mef2c-AHF-Cre* | 18.5 | H&E | None |
| 20 | *Mef2c-AHF-Cre* | 18.5 | H&E | None |
| 21 | *Mef2c-AHF-Cre* | 18.5 | H&E | Aberrant RScA |
| **Summary** | | | | **1/21 (4.8%)** |

**Supplemental Table 6. Cardiovascular defects in *VEGFR2*$^{Isl1\text{-}Het}$ and *VEGFR2*$^{SHF\text{-}Het}$ embryos. Abbreviations**: H&E – hematoxylin and eosin; µCT – micro-computed tomography; RScA – right subclavian artery.

### Table S7

| Cross | | Embryo numbers and genotypes | | | | | |
|---|---|---|---|---|---|---|---|
| | | **$VEGFR2^{SHF\text{-}WT}$** | **$VEGFR2^{SHF\text{-}Het}$** | **$VEGFR2^{SHF\text{-}KO}$** | **Total** | **χ2** | **p-value** |
| ***VEGFR2$^{+/-}$;mef2c-AHF-Cre x VEGFR2$^{flox/flox}$*** | Observed | 26 (40.6%) | 22 (34.4%) | 16 (25%) | 64 | 2.375 | 0.3050 |
| | Expected | 21.3 (33.3%) | 21.3 (33.3%) | 21.3 (33.3%) | | | |
| | | **$VEGFR2^{Isl1\text{-}WT}$** | **$VEGFR2^{Isl1\text{-}Het}$** | **$VEGFR2^{Isl1\text{-}KO}$** | **Total** | **χ2** | **p-value** |
| ***VEGFR2$^{+/-}$;Isl1$^{Cre/+}$ x VEGFR2$^{flox/flox}$*** | Observed | 21 (37.5%) | 17 (30.4%) | 18 (32.1%) | 56 | 0.4643 | 0.7928 |
| | Expected | 18.67 (33.3%) | 18.67 (33.3%) | 18.67 (33.3%) | | | |

**Supplemental Table 7. Loss of VEGFR2 in the SHF is not embryonic lethal.** Observed and expected numbers of embryos dissected between E14.5 and P0 for each genotype. Statistics were evaluated using the Chi-squared test.

### Table S8

| Genotype | Pharyngeal Arch | Number of ECs | Number of SHF-derived ECs | Number of Non-SHF-derived ECs |
|---|---|---|---|---|
| **Control** *Mef2c-AHF-Cre* (n = 6) | 4th | 434 ± 92 | 298 ± 74 | 136 ± 42 |
| **Tbx1$^{+/-}$** *Tbx1$^{+/-}$;Mef2c-AHF-Cre* (n = 6) | 4th | 330 ± 39 | 181 ± 16 | 149 ± 44 |
| **Tbx1$^{SHF\text{-}Het}$** *Tbx1$^{flox/+}$;Mef2c-AHF-Cre* (n = 6) | 4th | 483 ± 96 | 313 ± 67 | 170 ± 54 |
| **Tbx1$^{SHF\text{-}Null}$** *Tbx1$^{flox/flox}$;Mef2c-AHF-Cre* (n = 6) | 4th | 465 ± 85 | 174 ± 53 | 291 ± 51 |

**Supplemental Table 8. The average number of ECs in the pharyngeal arches of Tbx1 controls and mutants.** Pharyngeal arches of E10.5 control, *Tbx1$^{+/-}$*, *Tbx1$^{SHF\text{-}Het}$*, and *Tbx1$^{SHF\text{-}Null}$* embryos ranging from 34-36s were analyzed via WM-IF staining to detect VEGFR2, ERG, and Cre-reporter. The total number of ECs was determined in each 4th arch, and the percent SHF- and non-SHF-derived ECs were calculated. Averages ± SD are shown. Statistical analyses are reported in **Fig. 5D-F**.
