## Supplementary material for "Identification of novel buffering mechanisms in aortic arch artery development and congenital heart disease": Methods

**MATERIALS AND METHODS**

**Mice.** The mice were housed in Individually Ventilated Cage (IVC) racks (Allentown NexGen). Food and water were provided *ad libitum*, food was available through a feeder in the cage and water from the automatic watering system. The food is 5053 Purina PicoLab Rodent 20 for the non-mating cages. The mating cages were fed with 5058 Purina PicoLab Mouse 20. The light cycle is 12 hours on and 12 hours off, where the lights went on at 7 AM and off at 7 PM. The room temperature set point is 72^o^F. The following purchased strains were used in this study: *C57BL/6J* (The Jackson Laboratory, stock 0664); *Tg(Mef2c-AHF-Cre)2Blk/Mmnc* (MMRRCC, stock 030262- UNC) ^1^; *Kdr^tm2.1Jrt/J^* (The Jackson Laboratory, stock 017006) ^2^; *Kdr^tm2Sato/J^* (The Jackson Laboratory, stock 018977) ^3^; *B6.Cg-Gt(ROSA)26Sor^tm9(CAG-tdTomato)Hze/J^* from here on referred to as *Rosa^tdTomato^* Cre-reporter (The Jackson Laboratory, stock 007909) ^4^; and *B6.129(Cg)-Gt(ROSA)26Sor^tm4(ACTB-tdTomato,-EGFP)Luo/J^* from here on referred to as *Rosa^mTmG^* Cre-reporter (The Jackson Laboratory, stock 007676) ^5^. The following strains were gifted: *Isl1^Cre/+^* ^6^, *Tbx1^+/-^* ^7^, Tbx1^f/f^; RCE:loxP/RCE:loxP ^8,9^, where RCE:loxP allele is a Cre-reporter in the ROSA locus encoding enhanced green fluorescent protein (eGFP), *Apj-CreERT2* ^10^ and *Dre-Reporter* ^11^. Cre and Dre alleles were always transmitted through the male germline in all crosses. This is especially important for the Mef2c-AHF-Cre lines because when the Mef2c-AHF-Cre transgene is transferred through the female germline, it acts in all cells ^12,13^. The fidelity of cell-type specific Cre- or Dre-mediated recombination was assayed by the expression of fluorescent reporters. All previously published alleles were genotyped according to the protocols cited in the publications above. The sex of embryos was determined by PCR amplification of the *SRY* gene segment on the Y-chromosome, as described in ^14^. Equal numbers of male and female embryos were analyzed.

*Apj^T2A-DreERT2/+^* knock-in mice, hereafter referred to as *Apj^DreERT2/+^*, were generated by Biocytogen in C57BL/6J genetic background using CRISPR/Cas9-mediated mutagenesis and 5’- GTCCCAGCCCTAGTCCACAA -3’ gRNA sequence. The sequence encoding T2A-DreERT2 ^15,16^ was inserted using homologous recombination, replacing the TAA termination codon at the 3’end of the *Aplnr* gene. Mice containing correctly targeted *Aplnr* locus were selected following Southern blotting and Sanger sequencing. Sanger sequencing of the 10 top-predicted off-target sites confirmed the absence of mutations at these sites. The following primers were used for genotyping *Apj^T2A-DreERT2/+^*: wildtype forward 5’-TCTTTGACCCCCGATTTCGCCAAG-3’, wildtype reverse 5’-CAAGGCTCCATCCCTTT CCGCTAAA-3’, and Dre forward 5’-TTCTGTCCAGCACCCTGAAGTCTCT-3’. The wildtype allele gives a 392bp band, and the Dre allele gives a 596bp band. PCR conditions were 94°C for 5 min for one cycle, followed by 94°C for 30 sec, 60°C for 30 sec, and 72°C for 1 min for 35 cycles, and ending with 72°C for 5 min for one cycle.

*Kdr^tm2.1Jrt/J^* female mice, from here on referred to as *VEGFR2^+/-^*, were crossed with either *Tg(Mef2c-AHF-Cre)2Blk/Mmnc* or *Isl1^Cre/+^* male mice to generate *VEGFR2^+/-^;Mef2c-AHF-Cre* or *VEGFR2^+/-^; Isl1^Cre/+^* mice respectively. *Kdr^tm2Sato/J^*, from here on referred to as *VEGFR2^f/f^*, were crossed with either *Rosa^tdTomato^*, *Rosa^mTmG^*, or *Dre-Reporter* mice to generate homozygous *VEGFR2^f/f^*; *Rosa^tdTom/tdTom^*, *VEGFR2^f/f^*;*Rosa^mTmG/mTmG^*, and *VEGFR2^f/f^*;*Dre-Reporter/Dre-reporter* mice respectively. *VEGFR2^f/f^*;*Dre-Reporter* and *Apj^DreERT2/+^* mice were crossed to generate *VEGFR2^f/f^*; *Dre-Reporter; Apj^DreERT2/+^* mice. *Tbx1^+/-^* female mice were crossed with *Tg(Mef2c-AHF-Cre)2Blk/Mmnc* males to generate the strain referred to as *Tbx1^+/-^; Mef2c-AHF-Cre*. Tbx1^f/f^; RCE:loxP/RCE:loxP females were crossed with *Tg(Mef2c-AHF-Cre)2Blk/Mmnc* males to generate Tbx1f^/+-^;Mef2c-AHF-Cre male mice used for timed pregnancies.

*VEGFR2^GFP/+^; Mef2c-AHF-Cre*, *VEGFR2^GFP/+^; Isl1^Cre/+^*, *and Tbx1^+/-^ ; Mef2c-AHF-Cre, and Dre+* strains were maintained on *C57BL/6J* background for at least 10 generations. All other strains were maintained on a mixed background.

**Embryo collection and experimental design**

To collect embryos, females bearing two reporter alleles and the appropriate mutant alleles were crossed with males bearing the Cre or Dre sequences. The day of the vaginal plug was considered embryonic day (E) 0.5. Embryos were harvested on the desired day of development and fixed in 4% paraformaldehyde (PFA) overnight at 4°C. 4% PFA was prepared by diluting 16% PFA (Fisher, cat 50980487) and stored at -80^o^C until the day of use. Following fixation, embryos were washed in 1x Phosphate buffered saline (PBS, ThermoFisher, J75889-K2). Somites (s) were counted to stage embryos collected between E9.0-E10.5. Embryos were genotyped following previously established protocols. Each set of experiments was repeated at least three independent times. Each set of control, heterozygote, and mutant embryos were stained together using the same solutions to avoid differences due to technical variability in the solution preparation. The number (n) of biological replicates is stated in each figure legend.

**Tamoxifen Injections.** Pregnant females were injected intraperitoneally with 1.5 mg of tamoxifen (Sigma, T5648) solution dissolved in sesame oil (Sigma, S3547) on day E8.5 or E9.5, as indicated in figure legends. Embryos were dissected on day E10.5, fixed, stained, and imaged, as described below.

All experiments using mice were approved by the Institutional Animal Care and Use Committee of Rutgers University.

**Whole-mount Immunofluorescence.** E10.5 embryos were stained via whole-mount immunofluorescence according to the protocol previously described by Ramirez and Astrof ^17^. In brief, after overnight fixation at 4^o^C, embryos were washed with phosphate buffered saline (1x PBS prepared from 10xPBS solution, VWR, cat # 97063-660), then permeabilized with 0.1% Triton-X-100 in 1x PBS (PBST) overnight at 4^o^C, and blocked using 10% of normal donkey serum (Sigma, cat # D9663) in PBST, overnight at 4^o^C and then incubated with primary antibodies diluted in the blocking buffer for 4 days. After washing, embryos were incubated with secondary antibodies and DAPI diluted in the blocking buffer for 4 days. Following extensive washing, embryos were positioned in agarose, dehydrated through methanol series and cleared in benzyl alcohol/ benzyl benzoate (BABB). Cleared samples were mounted in BABB between two coverslips (VWR, cat # 16004-312) separated by a rubber spacer (Grace Bio Labs, cat # 664113), and imaged using confocal microscopy. Staining controls included: 1) incubations with secondary antibodies and 2) GFP-negative and mCherry-negative tissues to assess specificity of anti-GFP and anti-mCherry antibodies. To quantify the total number of ECs and SHF-derived ECs in the pharyngeal arches, embryos were harvested at E10.5, and somite staged to select embryos at the 34-37s stages; for quantification of venous ECs in the PAs, embryos were harvested at E9.5-10.0 and somite staged to select embryos at the 27-30s stages. The following 1° antibodies were used: anti-ERG (Abcam ab214341, 1:1000 or Abcam ab92513, 1:1000), anti-VEGFR2 (R&D Systems, AF644, 1:200), anti-Pecam 1 (BD Pharmingen, 553370, 1:200), anti-mCherry (Abcam ab167453, 1:1000), Sox17 (Abcam, ab224637, 1:100), and anti-COUPTFII (R&D Systems, PP-H714700, 1:1000). Alexa Fluor-conjugated 2° antibodies were used at 1:300 dilution and were purchased from ThermoFisher or Jackson Immunoresearch (see Antibodies table at the end of the methods section). DAPI (ThermoFisher, D3571, 1:1000 dilution of 5mg/ml solution) was used for staining nuclei. Control and mutant embryos were stained in pairs using the same antibody solutions.

**Whole-mount single-molecule RNA fluorescent *in situ* hybridization.** To visualize mRNA expression in embryos, whole mount *in situ* hybridization was performed according to the protocol described in Nomaru, Liu, De Bono, Righelli, Cirino, Wang, Song, Racedo, Dantas, Zhang, Cai, Angelini, Christiaen, Kelly, Baldini, Zheng and Morrow ^18,^Phillips, Stothard, Shaikh Qureshi, Kousa, Briones-Leon, Khasawneh, O'Loughlin, Sanders, Mazzotta, Dodds, Seidel, Bates, Nakatomi, Cockell, Schneider, Mohun, Maehr, Kist, Peters and Bamforth ^19^. Embryos were harvested at E9.5 and somite-staged to select embryos at 18-24s or 27-30s. Embryos were permeabilized using Protease III (Advanced Cell Diagnostics, 322381) for 20 minutes at room temperature. The following probes used were purchased from Advance Cell Diagnostics: Mm-Aplnr (436171) and Mm-Vegfa (405131-C2). The Vegfa-C2 probe was diluted (1:50) according to the manufacturer’s protocol. Amplification was done using RNAscope^®^ Multiplex Fluorescent v2 reagents (Advanced Cell Diagnostics, 323100). mRNA was visualized using TSA^®^ Plus Cyanine 3 (Akoya Biosciences, NEL744001KT) and Cyanine 5 (Akoya Biosciences, NEL745001KT) detection kits. Following *in situ* hybridization, embryos were stained to detect the VEGFR2 protein, as described above.

**Imaging.** E10.5 embryos were embedded in 1% agarose and cleared in 1:2 Benzyl Alcohol: Benzyl Benzoate (v/v), as previously described in Ramirez and Astrof ^17^. Confocal imaging was performed using a Nikon A1R microscope with 20x CFI Apo LWD Lambda S water immersion objective (MRD77200) or 25x CFI Plan Apo Lambda S silicone oil objectives (MRD73250) and NIS-Elements AR 5.11.01 64-bit software.

**Pharyngeal Arch Artery Analysis.**

a. ***Cell quantifications.*** Imaris software version 9.2.0 (Bitplane, USA) was used to segment pharyngeal arches, and the spot function was used for cell quantification, following the procedure outlined in Ramirez and Astrof ^17^. To determine the total number of ECs, ERG+ cells were quantified within each arch. To determine the total number of SHF-derived cells, Cre-reporter+ cells were quantified within each arch. The total number of SHF-derived ECs was determined by quantifying the number of ERG+, Cre-reporter+ ECs within each arch. Percent of SHF-derived ECs within each arch was calculated by dividing the number of ERG+, Cre-reporter+ cells by the total number of ERG+ cells.
b. ***Contribution of venous-derived cells to the 4^th^ PAA.*** Imaris software version 9.2.0 (Bitplane, USA) was used for cell quantifications. To determine the number of venous ECs within the 4th pharyngeal arch, the number of COUPTFII+, VEGFR2+ double-positive cells or the number of *Aplnr*+, VEGFR2+ double-positive cells was quantified. To determine Dre-recombinase labeling efficiency, we quantified the percentage of Dre-reporter+ ECs in the cardinal vein (**Fig. S3F, G**). Venous contribution to the 4^th^ PAA was calculated by normalizing the percent of Dre-reporter-labeled PAA ECs by Dre labeling efficiency.
c. ***Proliferation.*** Cell proliferation was assayed using EdU incorporation. EdU was diluted in saline (0.9% NaCl in water) to make a stock solution of 2.5 mg/ml. Pregnant females were injected with 250 μl of 0.5 μg/μl of EdU solution 30 min prior to dissection at E10.5. Embryos were fixed and stained with antibodies to VEGFR2 and tdTomato as above, and EdU was detected by incubating embryos in the EdU reaction cocktail (Click-iT EdU Alexa Fluor 555 Imaging Kit, cat # C10338) for 2 days at 4^o^C on a shaker in the dark. Nuclei were detected using DAPI. Embryos were then washed for 2 days in 1x PBST, changing 1xPBST solution intermittently. Following the washes, embryos were dehydrated using graded methanol series and cleared in BABB, as described above, prior to imaging using confocal microscopy. Imaris software was used to segment pharyngeal arches and the endothelium and to quantify the total number of endothelial nuclei and the number of endothelial nuclei that have incorporated EdU.
d. ***Mean intensity quantifications.*** To determine the level of *Vegfa* mRNA expression between control and mutant embryos, 0.5 μm optical sagittal sections spanning the pharyngeal apparatus were analyzed using Fiji ^20^. mRNA expression was quantified by measuring *Vegfa* mRNA signal intensity in the pharyngeal endoderm and SHF. The mean intensity of these regions was normalized to the mean intensity of the *Vegfa* mRNA signal in the dorsal side of the embryo (unaffected by the deletion) in the same image. 5-20 sections spaced every 5-10 μm were analyzed per side per embryo. Data were plotted as SuperPlots, wherein small dots indicated technical replicates, in this case each technical replicate was one optical section; Means for each biological replicate were marked by large dots of the same color as small dots; Only mean values for each biological sample were used for statistical calculations, as described in ^21^

**Analysis of PAA-derived vasculature, the aortic arch, its branches, and ductus.** Embryos were harvested on either day E14.5 or E18.5 and fixed with 4% PFA at room temperature following incisions in the abdomen and crown to allow for perfusion of PFA. Embryos were then rinsed with 1x PBS.

a. ***Hematoxylin and eosin (H&E).*** Samples were decalcified using formic acid-sodium citrate solution according to the protocol described in Morse ^22^. Formic acid-sodium citrate was made by combining two solutions, A and B, in equal parts. Solution A comprises 1:1 90% formic acid (Sigma, F0507) and distilled H_2_O (v/v). Solution B includes 20g of sodium citrate (Sigma, C8532) dissolved in 100 ml of distilled H_2_O. Embryos were then washed in H_2_O, followed by washes in graded series of ethanol (70%, 80%, 90%, and 100%), incubating embryos for 24 hours in each solution. Samples were then incubated in xylene for two days, changing the xylene solution each day, followed by embedding in paraffin. Paraffin blocks were sectioned at 6 μm and stained with H&E.
b. ***Micro-computed tomography (μCT)*** was performed using E18.5 embryos following methods and imaging described in Degenhardt, Wright, Horng, Padmanabhan and Epstein ^23^. All images were converted into 3-dimensional (3D) datasets using Imaris File Converter (version 9.2.0). Cardiovascular structures were manually segmented using Imaris software (Bitplane, USA).
c. ***Immunofluorescence on frozen sections.*** E14.5 embryos were dissected and fixed in 4% PFA at 4^o^C overnight with their abdomens open. Following fixation, embryos were washed in 1 x PBS and cryopreserved using incubations in increasing concentrations of sucrose (Fisher, 57-50-1) in 1 x PBS*: I)* 10% sucrose, rotating for 4 hours (hr) at room temperature (rt), *II)* 20% sucrose, rotating overnight at 4^o^C, and *III)* 30% sucrose, rotating 3 – 4 hr at rt. Embryos were then incubated in 1:1 mixture of 30% sucrose and OCT (Tissue-Tek, 4583), rotating 1 hr at rt and then placed into 100% OCT for 20 min without shaking until they sunk to the bottom and then positioned for cutting in the transverse orientation in the OCT inside cryomolds (VWR, 15160-099). Freezing was done by immersing the bottom of the molds into a beaker of 2-methylbutane (Fisher, O3551-4) pre-chilled on dry ice. Frozen blocks were stored at -80^o^C until use. 5-μm sections were collected on SuperFrost® PLUS Adhesion Slides (Electron Microscopy Sciences, 100501-220). Prior to staining, OCT was removed by washing the slides in 1X PBS. Barriers between sections were drawn using ImmEdge Hydrophobic barrier Pen (Vector Labs, H-4000). Sections were blocked using a blocking buffer containing 10% Donkey serum in PBST and incubated with primary antibodies anti-Pecam-1 (BD Pharmingen, 553370, 1:200) and anti-Sox17 (Abcam, ab224637, 1:100) in blocking solution at 4^o^C overnight. Slides were washed 3 x 10 min in PBST at rt, and then incubated in secondary antibodies and DAPI for 1 hr at rt. Sections were washed, coversliped using 1:1 mixture of 1 x PBS and glycerol as a mounting medium. Sections were imaged using confocal microscopy. The expression of mCherry (reporter) and GFP from the Flk1-GFP knockin ^24^ were detected by imaging their native fluorescence using 561 nm and 488 nm lasers.
d. **For whole-mount staining and imaging of the aortic arch, its branches, and the ductus arteriosus**, E14.5 embryos were fixed in 4% PFA in PBS for 24 hours at 4^o^C, washed in PBS, and then sectioned using Leica VT1000S vibratome to generate 900-micron-thick slices. Slices were stained according to the same procedure as E10.5 embryos in 24-well plates and cleared in BABB, as described in ^17^. Slices were imaged without embedding them into agarose, as described above.

**Statistics.** All statistics were performed using GraphPad Prism version 10 for Mac OS, GraphPad Software, San Diego, California, USA. Data were analyzed using non-parametric tests and the results were corrected for multiple testing. Tests used for specific experiments and figure panels are described in figure legends. Test of proportions for data in **Figure 2I** was performed using two different tests, both showed statistically significant results: 1) Log Odd Ratios with Woolf’s method for constructing confidence intervals; 2) Difference in Proportion test. In the tables below, WT, Het, or KO designate *VEGFR2^SHF-WT^, VEGFR2^SHF-Het^,* and *VEGFR2^SHF-KO^, respectively.* The explanation od statistical tests is provided in the Supplementary Statistical Analysis file.

| Log Odds ratio | Odds ratio | Std err | Arch | Z_score | P_val | Comparison | 99% CI Lower | 99% CI Upper |
| --- | --- | --- | --- | --- | --- | --- | --- | --- |
| 3.677 | 39.54 | 0.064 | 3 | 57.399 | 0 | WT vs. KO | 3.513 | 3.841 |
| 2.392 | 10.931 | 0.058 | 3 | 41.073 | 0 | Het vs. KO | 2.243 | 2.541 |
| 1.286 | 3.617 | 0.045 | 3 | 28.5 | 0 | WT vs. Het | 1.17 | 1.401 |
| 1.623 | 5.067 | 0.054 | 4 | 29.846 | 0 | WT vs. KO | 1.484 | 1.762 |
| 0.913 | 2.491 | 0.054 | 4 | 16.965 | 0 | Het vs. KO | 0.775 | 1.05 |
| 0.71 | 2.034 | 0.046 | 4 | 15.497 | 0 | WT vs. Het | 0.593 | 0.828 |
| 4.123 | 61.72 | 0.101 | 6 | 40.941 | 0 | WT vs. KO | 3.865 | 4.38 |
| 3.007 | 20.23 | 0.099 | 6 | 30.438 | 0 | Het vs. KO | 2.754 | 3.26 |
| 1.115 | 3.051 | 0.047 | 6 | 23.52 | 0 | WT vs. Het | 0.994 | 1.237 |

| Difference in Proportion | Std err | Arch | Z_score | P_val | Comparison | 99 % CI Lower | 99% Cl Upper |
| --- | --- | --- | --- | --- | --- | --- | --- |
| 0.658 | 0.008 | 3 | 81.618 | 0 | WT vs. KO | 0.637 | 0.679 |
| 0.353 | 0.007 | 3 | 51.131 | 0 | Het vs. KO | 0.336 | 0.371 |
| 0.305 | 0.01 | 3 | 31.259 | 0 | WT vs. Het | 0.28 | 0.33 |
| 0.263 | 0.009 | 4 | 30.628 | 0 | WT vs. KO | 0.241 | 0.285 |
| 0.118 | 0.007 | 4 | 17.506 | 0 | Het vs. KO | 0.101 | 0.136 |
| 0.145 | 0.009 | 4 | 15.412 | 0 | WT vs. Het | 0.121 | 0.169 |
| 0.637 | 0.009 | 6 | 74.569 | 0 | WT vs. KO | 0.615 | 0.658 |
| 0.366 | 0.008 | 6 | 46.514 | 0 | Het vs. KO | 0.346 | 0.386 |
| 0.271 | 0.011 | 6 | 24.883 | 0 | WT vs. Het | 0.243 | 0.298 |

**SUPPLEMENTARY MATERIALS AND METHODS**

**Mice**

| **Strain** | **Abbreviated Name** | **Vendor or Source** | **Background Strain** | **Other Information** | **Reference** |
| --- | --- | --- | --- | --- | --- |
| C57BL/6J | *B6WT* | The Jackson Laboratory, stock 0664 | C57BL/6J |  |  |
| Tg(Mef2c-AHF-Cre)2Blk/Mmnc | *Mef2c-AHF-Cre* | MMRRCC, stock 030262- UNC | C57BL/6J |  | ^1^ |
| *Kdr^tm2.1Jrt/J^* | *VEGFR2^GFP/+^* | The Jackson Laboratory, stock 017006 | C57BL/6J |  | ^2^ |
| *Kdr^tm2Sato/J^* | *VEGFR2^f/f^* | The Jackson Laboratory, stock 018977 | Mixed |  | ^3^ |
| *B6.Cg-Gt(ROSA)26Sor^tm9(CAG-tdTomato)Hze/J^* | *Rosa^tdTomato^ Cre-reporter* | The Jackson Laboratory, stock 007909 | Mixed |  | ^4^ |
| *B6.129(Cg)-Gt(ROSA)26Sor^tm4(ACTB-tdTomato,-EGFP)Luo/J^* | *Rosa^mTmG^ Cre-reporter* | The Jackson Laboratory, stock 007676 | C57BL/6J |  | ^5^ |
| B6;129S- Gt(ROSA)26Sor^tm66.1(CAG- tdTomato)Hze/J^ | *Ai66/Dre-reporter* | Gifted by Dr. Hongkui Zeng | 129/C57BL/6J | LoxP sites in this strain were recombined converting this strain to a Dre reporter strain. | ^11^ |
| *Isl1^tm1(cre)Sev^* | *Isl1^Cre/+^* | Gifted by Dr. Sylvia Evans | C57BL/6J |  | ^6^ |
| *Tbx1^+/-^* |  | Gifted by Dr. Bernice Morrow | C57BL/6J |  | ^7^ |
| *Tbx1^flox/flox^* |  | Gifted by Dr. Bernice Morrow | Mixed background |  | ^25^ |
| Aplnr-creER (Tg(Aplnr-cre/ERT2)^#Krh^) | *Apj-CreERT2* | Gifted by Dr. Kristy Red-Horse | C57BL/6J |  | ^10^ |
| Apj^T2A-DreERT2/+^ | *Apj^DreERT2/+^* | This study | C57BL/6J |  |  |

**Antibodies**

| **Target Antigen** | **Vendor/Source** | **Catalog #** | **Dilution Factor** |
| --- | --- | --- | --- |
| ERG | Abcam | ab92513 | 1:1000 |
| ERG | Abcam | ab214341 | 1:1000 |
| Cherry | Abcam | ab167453 | 1:1000 |
| COUP-TFII | R&D Systems | PP-H714700 | 1:1000 |
| Green fluorescent protein | Aves | GFP-1020 | 1:500 |
| Pecam-1 | BD Pharmingen | 553370 | 1:200 |
| VEGFR2 | R&D Systems | AF644 | 1:200 |
| Sox17 | Abcam |  |  |
| Alexa Fluor 488 donkey anti-goat | Thermofisher | A11055 | 1:300 |
| Alexa Fluor 488 donkey anti-rabbit | Thermofisher | A21206 | 1:300 |
| Alexa Fluor 555 donkey anti-rabbit | Thermofisher | A31572 | 1:300 |
| Alexa Fluor 555 donkey anti-mouse | Thermofisher | A31570 | 1:300 |
| Alexa Fluor 647 donkey anti-mouse | Thermofisher | A31571 | 1:300 |
| Alexa Fluor 647 donkey anti-goat | Thermofisher | A21447 | 1:300 |
| Alexa Fluor 488 donkey anti-chicken | Jackson Immunoresearch | 703-545-155 | 1:300 |

**ISH Probes**

| **Probe** | **Vendor/Source** | **Channel** | **Catalog Number** |
| --- | --- | --- | --- |
| Mm-AplnR | Advanced Cell Diagnostics | C1 | 436171 |
| Mm-Vegfa | Advanced Cell Diagnostics | C2 | 405131-C2 |

**Chemical/Kits**

| **Name** | **Vendor/Source** | **Catalog #** |
| --- | --- | --- |
| DAPI | Thermofisher | D3571 |
| Tamoxifen | Sigma | T5648 |
| Sesame oil | Sigma | S3547 |
| Protease 3 | Advance Cell Diagnostics | 322381 |
| RNAscope ® Multiplex Fluorescent v2 | Advance Cell Diagnostics | 323100 |
| TSA® Plus Cyanine 3 (Cy3) detection kit | Akoya Biosciences | NEL744001KT |
| TSA® Plus Cyanine 5 (Cy5) detection kit | Akoya Biosciences | NEL745001KT |
| Benzyl Alcohol | Sigma | 305197 |
| Benzyl Benzoate | Sigma | [B6630](https://b2b.sigmaaldrich.com/US/en/product/sial/b6630) |
| Formic Acid | Sigma | F0507 |
| Sodium Citrate | Sigma | C8532 |

1.	Verzi MP, McCulley DJ, De Val S, Dodou E, Black BL. The right ventricle, outflow tract, and ventricular septum comprise a restricted expression domain within the secondary/anterior heart field. In: *Developmental Biology*. 2005:134-145.

2.	Ema M, Takahashi S, Rossant J. Deletion of the selection cassette, but not cis-acting elements, in targeted Flk1-lacZ allele reveals Flk1 expression in multipotent mesodermal progenitors. In: *Blood*. 2006:111-117.

3.	Hooper AT, Butler JM, Nolan DJ, Kranz A, Iida K, Kobayashi M, Kopp HG, Shido K, Petit I, Yanger K, et al. Engraftment and Reconstitution of Hematopoiesis Is Dependent on VEGFR2-Mediated Regeneration of Sinusoidal Endothelial Cells. In: *Cell Stem Cell*. 2009:263-274.

4.	Madisen L, Zwingman TA, Sunkin SM, Oh SW, Zariwala HA, Gu H, Ng LL, Palmiter RD, Hawrylycz MJ, Jones AR, et al. A robust and high-throughput Cre reporting and characterization system for the whole mouse brain. *Nat Neurosci*. 2010;13:133-140. doi: 10.1038/nn.2467

5.	Muzumdar MD, Tasic B, Miyamichi K, Li L, Luo L. A global double-fluorescent Cre reporter mouse. *Genesis*. 2007;45:593-605. doi: 10.1002/dvg.20335

6.	Yang L, Cai CL, Lin L, Qyang Y, Chung C, Monteiro RM, Mummery CL, Fishman GI, Cogen A, Evans S. Isl1Cre reveals a common Bmp pathway in heart and limb development. *Development*. 2006;133:1575-1585. doi: 10.1242/dev.02322

7.	Merscher S, Funke B, Epstein JA, Heyer J, Puech A, Lu MM, Xavier RJ, Demay MB, Russell RG, Factor S, et al. TBX1 is responsible for cardiovascular defects in velo-cardio-facial/DiGeorge syndrome. In: *Cell*. 2001:619-629.

8.	Arnold SM, Fessler LI, Fessler JH, Kaufman RJ. Two homologues encoding human UDP-glucose:glycoprotein glucosyltransferase differ in mRNA expression and enzymatic activity. *Biochemistry*. 2000;39:2149-2163. doi: 10.1021/bi9916473

9.	Sousa VH, Miyoshi G, Hjerling-Leffler J, Karayannis T, Fishell G. Characterization of Nkx6-2-derived neocortical interneuron lineages. *Cereb Cortex*. 2009;19 Suppl 1:i1-10. doi: 10.1093/cercor/bhp038

10.	Chen HI, Sharma B, Akerberg BN, Numi HJ, Kivela R, Saharinen P, Aghajanian H, McKay AS, Bogard PE, Chang AH, et al. The sinus venosus contributes to coronary vasculature through VEGFC-stimulated angiogenesis. In: *Development*. 2014:4500-4512.

11.	Madisen L, Garner AR, Shimaoka D, Chuong AS, Klapoetke NC, Li L, van der Bourg A, Niino Y, Egolf L, Monetti C, et al. Transgenic Mice for Intersectional Targeting of Neural Sensors and Effectors with High Specificity and Performance. In: *Neuron*. 2015:942-958.

12.	Ehlers ML, Celona B, Black BL. NFATc1 controls skeletal muscle fiber type and is a negative regulator of MyoD activity. *Cell Rep*. 2014;8:1639-1648. doi: 10.1016/j.celrep.2014.08.035

13.	Sinha T, Lammerts van Bueren K, Dickel DE, Zlatanova I, Thomas R, Lizama CO, Xu SM, Zovein AC, Ikegami K, Moskowitz IP, et al. Differential Etv2 threshold requirement for endothelial and erythropoietic development. *Cell Rep*. 2022;39:110881. doi: 10.1016/j.celrep.2022.110881

14.	Astrof S, Kirby A, Lindblad-Toh K, Daly M, Hynes RO. Heart development in fibronectin-null mice is governed by a genetic modifier on chromosome four. *Mech Dev*. 2007;124:551-558. doi: 10.1016/j.mod.2007.05.004

15.	Devine WP, Wythe JD, George M, Koshiba-Takeuchi K, Bruneau BG. Early patterning and specification of cardiac progenitors in gastrulating mesoderm. In: *eLife*. 2014:1-23.

16.	Liu Z, Chen O, Wall JBJ, Zheng M, Zhou Y, Wang L, Vaseghi HR, Qian L, Liu J. Systematic comparison of 2A peptides for cloning multi-genes in a polycistronic vector. *Sci Rep*. 2017;7:2193. doi: 10.1038/s41598-017-02460-2

17.	Ramirez A, Astrof S. Visualization and Analysis of Pharyngeal Arch Arteries using Whole-mount Immunohistochemistry and 3D Reconstruction. *J Vis Exp*. 2020. doi: 10.3791/60797

18.	Nomaru H, Liu Y, De Bono C, Righelli D, Cirino A, Wang W, Song H, Racedo SE, Dantas AG, Zhang L, et al. Single cell multi-omic analysis identifies a Tbx1-dependent multilineage primed population in murine cardiopharyngeal mesoderm. *Nat Commun*. 2021;12:6645. doi: 10.1038/s41467-021-26966-6

19.	Phillips HM, Stothard CA, Shaikh Qureshi WM, Kousa AI, Briones-Leon JA, Khasawneh RR, O'Loughlin C, Sanders R, Mazzotta S, Dodds R, et al. Pax9 is required for cardiovascular development and interacts with Tbx1 in the pharyngeal endoderm to control 4th pharyngeal arch artery morphogenesis. In: *Development*. 2019:dev177618.

20.	Schindelin J, Arganda-Carreras I, Frise E, Kaynig V, Longair M, Pietzsch T, Preibisch S, Rueden C, Saalfeld S, Schmid B, et al. Fiji: an open-source platform for biological-image analysis. *Nat Methods*. 2012;9:676-682. doi: 10.1038/nmeth.2019

21.	Lord SJ, Velle KB, Mullins RD, Fritz-Laylin LK. SuperPlots: Communicating reproducibility and variability in cell biology. *J Cell Biol*. 2020;219. doi: 10.1083/jcb.202001064

22.	Morse A. Formic Acid-Sodium Citrate Decalcification and Butyl Alcohol Dehydration of Teeth and Bones for Sectioning in Paraffin. *Journal of Dental Research*. 1945:143-153. doi: 10.1177

23.	Degenhardt K, Wright AC, Horng D, Padmanabhan A, Epstein JA. Rapid 3D Phenotyping of Cardiovascular Development in Mouse Embryos by Micro-CT With Iodine Staining. In: *Circulation: Cardiovascular Imaging*. 2010:314-322.

24.	Ema M, Takahashi S, Rossant J. Deletion of the selection cassette, but not cis-acting elements, in targeted Flk1-lacZ allele reveals Flk1 expression in multipotent mesodermal progenitors. *Blood*. 2006;107:111-117. doi: 10.1182/blood-2005-05-1970

25.	Arnold JS, Werling U, Braunstein EM, Liao J, Nowotschin S, Edelmann W, Hebert JM, Morrow BE. Inactivation of Tbx1 in the pharyngeal endoderm results in 22q11DS malformations. *Development*. 2006;133:977-987. doi: 10.1242/dev.02264
