## Supplemental statistical analysis for "Identification of novel buffering mechanisms in aortic arch artery development and congenital heart disease"

Stratified Analysis

2024-02-01

### Statistical Setup

Suppose we have two genotypes A and B, and we wish to study how cells differentiate into the two categories [SHF-derived] and [non-SHF-derived]. The collected data can be represented by a $2\times2$ contingency as shown below. The cell counts are given by $n_{11},n_{12},n_{21}$, and $n_{22}$.

|  | SHF-derived | Non-SHF-derived |
| --- | --- | --- |
| Genotype A | $n_{11}$ | $n_{12}$ |
| Genotype B | $n_{21}$ | $n_{22}$ |

We can view the data as coming from two independent populations, with genotypes A and B represent the first and second populations respectively. Holding the row totals fixed, we can statistically model each row as coming from the independent binomial distribution:

$$n_{11}\sim\text{Bin}\left( n_{11}+n_{12},\pi_{a} \right)$$

$$n_{21}\sim\text{Bin}\left( n_{21}+n_{22},\pi_{b} \right).$$

Note, we need only specify a distribution for the first cell in each row since we have fixed row totals. An estimate for $\pi_{a}$ and $\pi_{b}$ can be found by using the sample proportion, which gives us the estimates:

$$\hat{\pi}_{a}=\frac{n_{11}}{n_{11}+n_{12}}$$

$$\hat{\pi}_{b}=\frac{n_{21}}{n_{21}+n_{22}}.$$

Critically, this statistical model assumes the two populations are independent. This is a reasonable assumption in our case since each embryo can only have one genotype. The question of whether or not the two genotypes are different corresponds to checking whether or not $\pi_{a}=\pi_{b}$, with equality holding if there is no difference. Thus a hypothesis test for checking whether or not genotype has an effect can be formulated where the null is $H_{0}:\pi_{a}-\pi_{b}=0$, and the alternative is $H_{1}:\pi_{a}-\pi_{b}\neq0$. The next two sections will go over two different ways to carry this hypothesis test.

### Difference of Proportions

An intuitive approach to testing the null is to directly estimate $\pi_{a}-\pi_{b}$ using the difference in sample proportions, $\hat{\pi}_{a}-\hat{\pi}_{b}$. When the row totals are large, the sampling distributions for $\hat{\pi}_{a}$ and $\hat{\pi}_{b}$ are well approximated by the normal distribution. More precisely, we have

$$\hat{\pi}_{a}\sim N\left( \pi_{a},\frac{\pi_{a}\left( 1-\pi_{a} \right)}{n_{11}+n_{12}} \right)$$

$$\hat{\pi}_{b}\sim N\left( \pi_{b},\frac{\pi_{b}\left( 1-\pi_{b} \right)}{n_{21}+n_{22}} \right).$$

Under the null hypothesis, the test statistic, $\hat{\pi}_{a}-\hat{\pi}_{b}$, approximately follows a normal distribution with mean and variance

$$\hat{\pi}_{a}-\hat{\pi}_{b}\sim N\left( 0,\frac{\pi_{a}\left( 1-\pi_{a} \right)}{n_{11}+n_{12}}+\frac{\pi_{b}\left( 1-\pi_{b} \right)}{n_{21}+n_{22}} \right).$$

Using the preceding approximation, we can calculate a $Z$-score and the associated $p$-value. One notable issue is that in order to calculate a $Z$-score we must divide by the standard error, which is a function of the unknown parameters $\pi_{a}$ and $\pi_{b}$. This can be easily fixed by replacing them with the corresponding estimates $\hat{\pi}_{a}$ and $\hat{\pi}_{b}$. This substitution requires a larger sample size to be accurate, but is not a concern in our case.

### Log Odds Ratio

A popular alternative test statistic to the sample proportion, is the sample log odds ratio. The odds ratio is another way of measuring the difference between $\pi_{a}$ and $\pi_{b}$. To introduce how the odds ratio is defined, we must first define the odds of an event. If $\pi$ is the probability of a cell being [SHF-derived], then we say the odds of a cell being SHF-derived is $\pi/\left( 1-\pi\right)$. The odds ratio is a ratio between two odds. It describes the strength of association between a treatment and outcome in a $2\times2$ table. In our case, the treatment is the genotype and the outcome is the type of cell ([SHF-derived] or [Non-SHF-derived]). Let $\theta$ denote the odds ratio, then $\theta$ is defined as

$$\theta:=\frac{\pi_{a}/\left( 1-\pi_{a} \right)}{\pi_{b}/\left( 1-\pi_{b} \right)}.$$

To see how the difference in proportion is related to the odds ratio, note $\pi_{a}=\pi_{b}$ if and only if $\text{ln}\left( \theta\right)=0$. Thus, the null hypothesis $H_{0}:\pi_{a}-\pi_{b}=0$ is equivalent to $H_{0}:\text{ln}\left( \theta\right)=0$. Similar to the previous method, an estimate for $\theta$ can be found by plugging in $\hat{\pi}_{a}$ and $\hat{\pi}_{b}$ for $\pi_{a}$ and $\pi_{b}$ respectively. This gives us the estimate

$$\hat{\theta}=\frac{\hat{\pi}_{a}/\left( 1-\hat{\pi}_{a} \right)}{\hat{\pi}_{b}/\left( 1-\hat{\pi}_{b} \right)}=\frac{n_{11}n_{22}}{n_{12}n_{21}}.$$

An application of the delta method will show that taking the natural log of $\hat{\theta}$ will result in an approximately normal distribution with mean and variance given by

$$\text{ln}\left( \hat{\theta} \right)\sim N\left( \text{ln}\left( \theta\right),\frac{1}{n_{11}}+\frac{1}{n_{12}}+\frac{1}{n_{21}}+\frac{1}{n_{22}} \right).$$

This approximation can be use to calculate the associated $Z$-scores and $p$-values. The log odds ratio may be preferable compared to the difference in proportion because of its connection to the logistic regression model. In fact, the estimator $\hat{\theta}$ has a one-to-one correspondence to the maximum likelihood estimate for a certain logistic regression model.

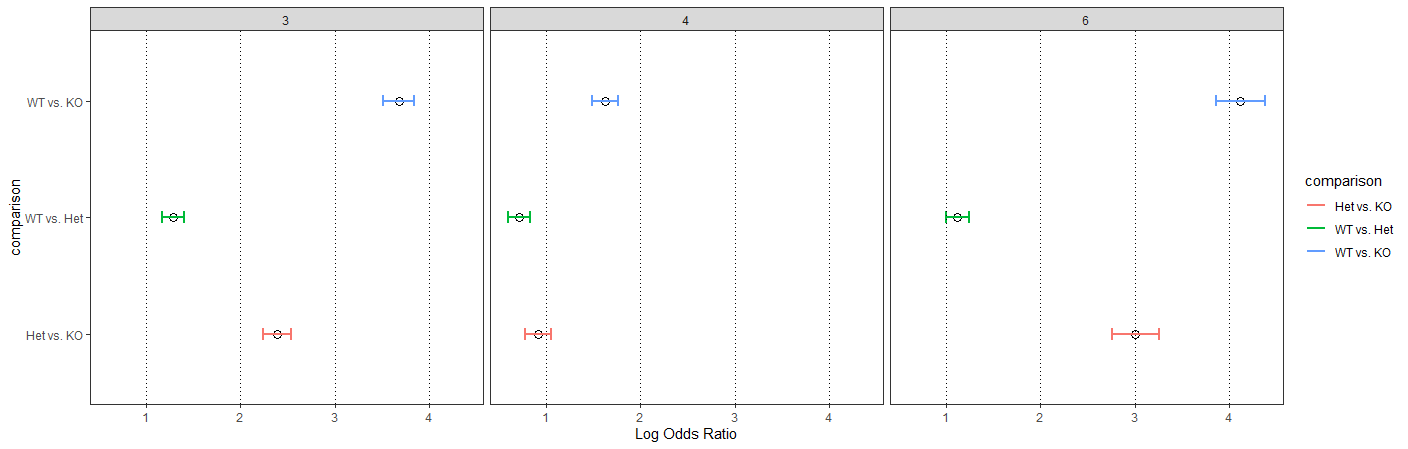

**Graphical representation of results of the Log Odds Ratio test for Figure 2I.** Error bars mark 99% confidence interval. Het – *VEGFR2^SHF-Het^*, KO – *VEGFR2^SHF-KO^*, WT – *VEGFR2^SHF-WT^.* 3, 4, and 6 – Pharyngeal arches 3, 4, and 6.
